# Magnesium and the magnesium transporter UEX regulate sleep via Ca^2+^-dependent CREB signaling and a CNK-ERK pathway

**DOI:** 10.1101/2022.09.26.509486

**Authors:** Xin Yuan, Huimei Zheng, Xiao Xu, Huan Deng, Xiaohang Yang, Yongmei Xi

## Abstract

Magnesium and its related preparations are already in medical use and have recognized therapeutic effects on sleep disorders. However, its underlying molecular mechanisms remain unclear. Here, using *Drosophila* as a model, we found that RNAi-mediated knockdown of *Uex*, the homologous gene of magnesium transporters of the Cyclin M family (CNNM) causes increased daily total sleep. Ectopic-expression of CNNM1 can rescue the sleep phenotype in *Uex* knockdown flies. UEX exhibits rhythmic oscillations in the brain and affects the efflux of cellular Mg^2+^. Knockdown of *Uex* in the nervous system influences Ca^2+^-mediated CREB signaling and neuroplasticity. Additionally, Uex physically interacts with CNK, the upstream regulator of ERK pathway. Similar effects on sleep are observed with knockdown of *Cnk* in flies. We propose that the UEX regulates sleep through its downstream Ca^2+^-dependent CREB signaling and a CNK-ERK pathway. Our findings may provide new insight into mechanisms of magnesium and magnesium transporter related sleep disorder.

## Introduction

The physiological regulation of sleep is related to many aspects such as metabolic balance, neurobehavioral control, and learning and memory (Girardeau and Lopes-Dos-Santos, 2021; Panda, 2016; Vyazovskiy et al., 2017). Many human diseases, especially neurological and/or aging-related diseases, are accompanied with sleep disorders(Borniger et al., 2018; Stefani and Hogl, 2020; Wulff et al., 2010). Magnesium is believed to play a role in calming and facilitating sleep and as an important element for maintaining the normal function of nerves and muscles. As such, magnesium preparations are often prescribed. Magnesium sulfate, for example, is used to treat migraine (Dolati et al., 2020), relieve sleep problems in patients with depression (Serefko et al., 2016), or help patients with preeclampsia to relax their nerves (Duley et al., 2010). Supplementation of magnesium can also help combat insomnia in the elderly (Abbasi et al., 2012; Nielsen et al., 2010). However, the mechanisms by which magnesium and magnesium transporters regulate sleep are unclear.

Ion transporters such as magnesium transporters are transmembrane proteins that actively transport ions across the cell membrane against their concentration gradients. It is a characteristically slow process which lies in direct contrast to ion channels through which much faster translocations can occur (Neverisky and Abbott, 2015). The active transport of Mg^2+^ through the cell membrane by a magnesium transporter is 10^3^ times slower than the passive transport of Ca^2+^ through its own channels (Hille, 2001). Functionally conserved magnesium transporters such as TRPM7 (Transient receptor potential cation channel subfamily M member 7), the CNNMs (cyclin M/ancient conserved domain proteins), SLC41A (Solute carrier family 41), MRS2 (Magnesium transporter MRS2), and MAGT1 (Magnesium transporter 1) are present from mammals to their homologs in flies (de Baaij et al., 2015; Yamazaki et al., 2013). It has been reported that the TRPM7 plays a role not in transporting magnesium but also in regulating calcium and zinc homeostasis (Georgiev et al., 2010; Hu and Wolfner, 2019). Among the CNNM protein family (CNNM1-4), CNNM4 was seen to act in regulating sperm Ca^2+^ homeostasis (Yamazaki et al., 2016), and has been reported linking to human diseases such as Jalili syndrome and hypomagnesemia (Parry et al., 2009; Stuiver et al., 2011). We have previously reported that *Drosophila Uex*, the homolog of the Mammalian CNNMs, interacts with PRL-1 to play a neuroprotective role in olfactory CO2 sensory circuitry (Guo et al., 2019).

In *Drosophila*, there are about 150 pacemaker neurons in each hemisphere of the adult brain, which can be distinguished into dorsal neurons (DNs) and lateral neurons (LNs) and defined by their related expressions of conserved rhythmic molecules such as Clock, PER, TIM, CRY and PDF (Gummadova et al., 2009; Guo et al., 2016; Peschel and Helfrich-Forster, 2011). These core pacemaker neurons are connected to many downstream neurons such as DH44 (Cavey et al., 2016), PDFR (Potdar et al., 2018), fan-shaped (Ni et al., 2019), ellipsoid body (Guo et al., 2018), par intercerebralis (PI) (Crocker et al., 2010; Park et al., 2014) or mushroom body (MB) neurons (Pitman et al., 2006). Combined with external environmental factors such as light, they form characteristic rhythms within the brain and act on behavioral output, thus affecting patterns of wakefulness or sleep (Alpert et al., 2020; Chen et al., 2018a).

In addition to these neurons that are accurately positioned in the circadian clock, many substances in the cells, potassium for example, show rhythmic fluctuations(Feeney et al., 2016; Flourakis et al., 2015). The potassium channel SHAKER affects *Drosophila* sleep, its mutation causing a short sleep phenotype (Cirelli et al., 2005; Kempf et al., 2019). The calcium channel TRPA1 can also monitor the ambient temperature and regulate changes in the sleep response of *Drosophila* to different temperatures (Roessingh and Stanewsky, 2017). Ca^2+^-permeable NMDAR channels, together with their downstream genes, also participate in sleep regulation (Liu et al., 2016; Lymer and Blau, 2016). Magnesium is an important ATP cofactor (Jahnen-Dechent and Ketteler, 2012), also acting as a regulator of voltage-dependent Ca^2+^ channels (Pilchova et al., 2017). Magnesium fluxes has been reported to regulate cellular timekeeping and energy balance (Feeney et al., 2016), which explains the rhythmic fluctuations of energy-dependent activities such as transcription and translation (Cao et al., 2011; de Baaij et al., 2015)).

In the present study, we identified that *Drosophila* UEX, as the homologous gene of CNNM, functions in regulating total sleep time. Our data show that UEX exhibits rhythmic oscillations at a protein level rather than mRNA level. Knockdown of *Uex* in the *Drosophila* nervous system affects Ca^2+^-dependent CREB signaling and neuroplasticity, resulting in a severe sleep phenotype including increased total daily sleep. Supplementation of Ca^2+^ could rescue the sleep phenotype in *Uex* knockdown flies. Furthermore, we found that UEX interacts with CNK and regulates phosphorylated ERK (ppERK) expression, and downregulation of CNK also induce an increase in total sleep time in flies. We proposed that magnesium and the magnesium transporter UEX regulate sleep via Ca^2+^-CREB signaling and CNK-ERK pathway.

## Results

### 1. Knockdown of *Uex* led to increased total daily sleep in *Drosophila*

We previously reported the neuroprotective role of the UEX/PRL-1 complex in *Drosophila* relating to olfactory CO_2_ stimulation (Guo et al., 2019). This indicated that *Uex* plays an important role in the nervous system. We further found that *Uex* knockdown driven by Elav-Gal4 (*Elav*>*Uex* RNAi) led to a sleep phenotype in flies that included loss of morning (Zeitgeber time 0, ZT0) and evening (ZT12) peaks (Figure 1A) and increased total daily sleep (Figure 1B) and sleep bout durations (Figure 1C), but with no obvious changes in the sleep bout number (Figure 1D). We analyzed the locomotor ability of *Elav*>*Uex* RNAi flies during their waking periods and found that this was unaffected (Figure 1E), remaining comparable to that of wild type flies. RNAi line alone have no obvious phenotype exclude the background effects. Additionally, western Blots were carried out to identify the knock down efficiency of *Uex*, revealing the protein levels of UEX in the head tissue as significantly downregulated, compared to controls (Figure 1–figure supplement 1A-1B). This formed a reliable foundation for the following experiments.

**Figure 1.**
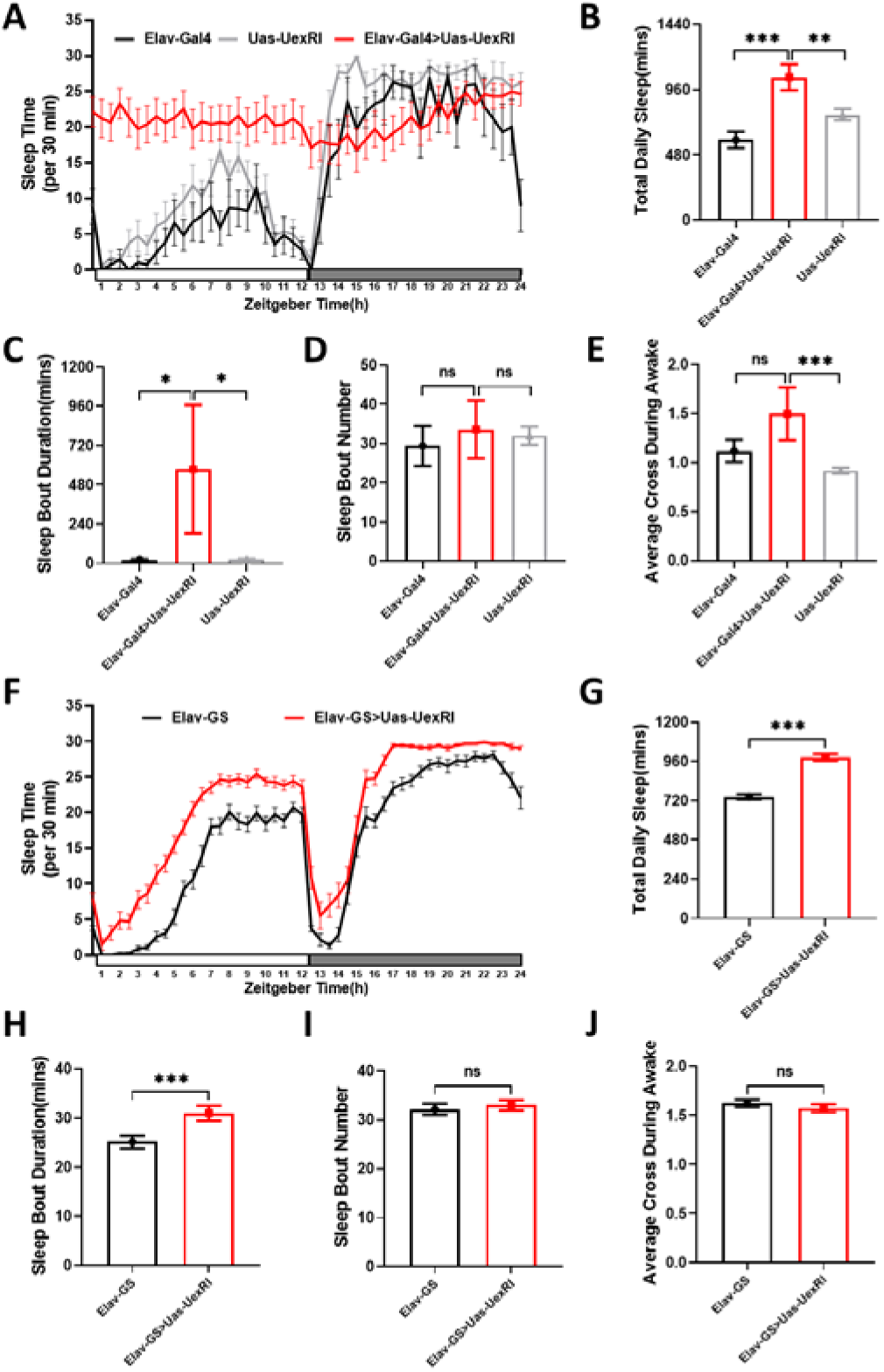
Knockdown of *Uex* led to increased total daily sleep in *Drosophila*. (A) Typical sleep profile of Elav-Gal4 flies (black line, n=16), UAS-Uex RNAi flies (gray line, n=16) and Elav-Gal4 > UAS-Uex RNAi flies (red line, n=11). Sleep time was plotted in 30-min bin. The x-coordinate represents zeitgeber time and the y-coordinate represents sleep time every 30 minutes. (B) The total daily sleep quantification in (A). (C) The mean sleep bout duration quantification in (A). (D) The number of quantified sleep bouts in (A). (E) The quantification of average cross counts per min in (A). (F) Typical sleep profile of Elav-GS flies (black line, n=32) and Elav-GS > UAS-Uex RNAi flies (red line, n=32). Sleep time was plotted in 30-min bin. The x-coordinate represents zeitgeber time and the y-coordinate represents sleep time every 30 minutes. (G) The total daily sleep quantification in (F). (H) The mean sleep bout duration quantification in (F). (I) The number of quantified sleep bouts in (F). (J) The quantification of average cross counts per min in (F).

Since the expression of pan-neuronal Elav-GAL4 begins from the larval stage (Yao and White, 1994), in addition to altered sleep behavior, *Elav*>*Uex* RNAi flies also show other alterations in behaviours not directly related to sleep such as displaying a permanent wing held-up phenotype (Guo et al., 2019). In order to exclude the effects of *Uex* knockdown on development, we utilized the gene-switch (GS) method to induce RNA interference only in adult fly neurons (Osterwalder et al., 2001). The expression of GAL4 protein is induced by the addition of the anti-progestin mifepristone (RU486) drug to the food. Results showed that RU486-induced *Uex* knockdown also led to increased total daily sleep and increased sleep bout durations (Figures 1F-1H), as was the case for *Elav*>*Uex* RNAi flies, with no obvious changes in the sleep bout number and locomotor abilities remaining unaffected (Figures 1J-1I).

### 2. UEX functions in multiple brain regions containing PDF, MB and PI neurons

To determine the specific type of neurons in which the UEX protein plays a role in regulating sleep behavior, we performed screening experiments in two ways. Firstly, we used various sleep-related GAL4 drivers to knockdown *Uex* in different brain regions, including neurotransmitter neurons, circadian neurons, and the specific neurons in interbrain area, mushroom body, and central complex regions, respectively. The data revealed that specific knockdown of *Uex*, in PDF neurons, MB neurons, PI neurons, compounded C309 neurons or 121Y neurons led to a significant increase in total daily sleep (Figure 2A and S1C). Notably, the knockdown of *Uex* in 121Y-GAL4 expressing neurons (121Y>*Uex* RNAi) resulted in the most severe phenotypes (Figures 2B), with increased total daily sleep and sleep bout duration (Figures 2C-2D), but reduced sleep bout number (Figure 2E). Immunostaining showed that the expression of 121Y-GAL4 covers multiple brain regions containing the PI neurons, MB neurons and PDF positive lateral ventral neurons (PDF^+^ LNvs) (Figure 2F). PI has been considered to serve as an essential circadian output region (Cavey et al., 2016). Notably, the knockdown of *Uex* in several PI labelling Gal4 drivers (including C929, DH44, 50Y, C309) all caused obvious daily sleep increases (Figures S1D-S1H), especially when conducted in DH44 sleep output neurons (Figures S1I-S1J).

**Figure 2.**
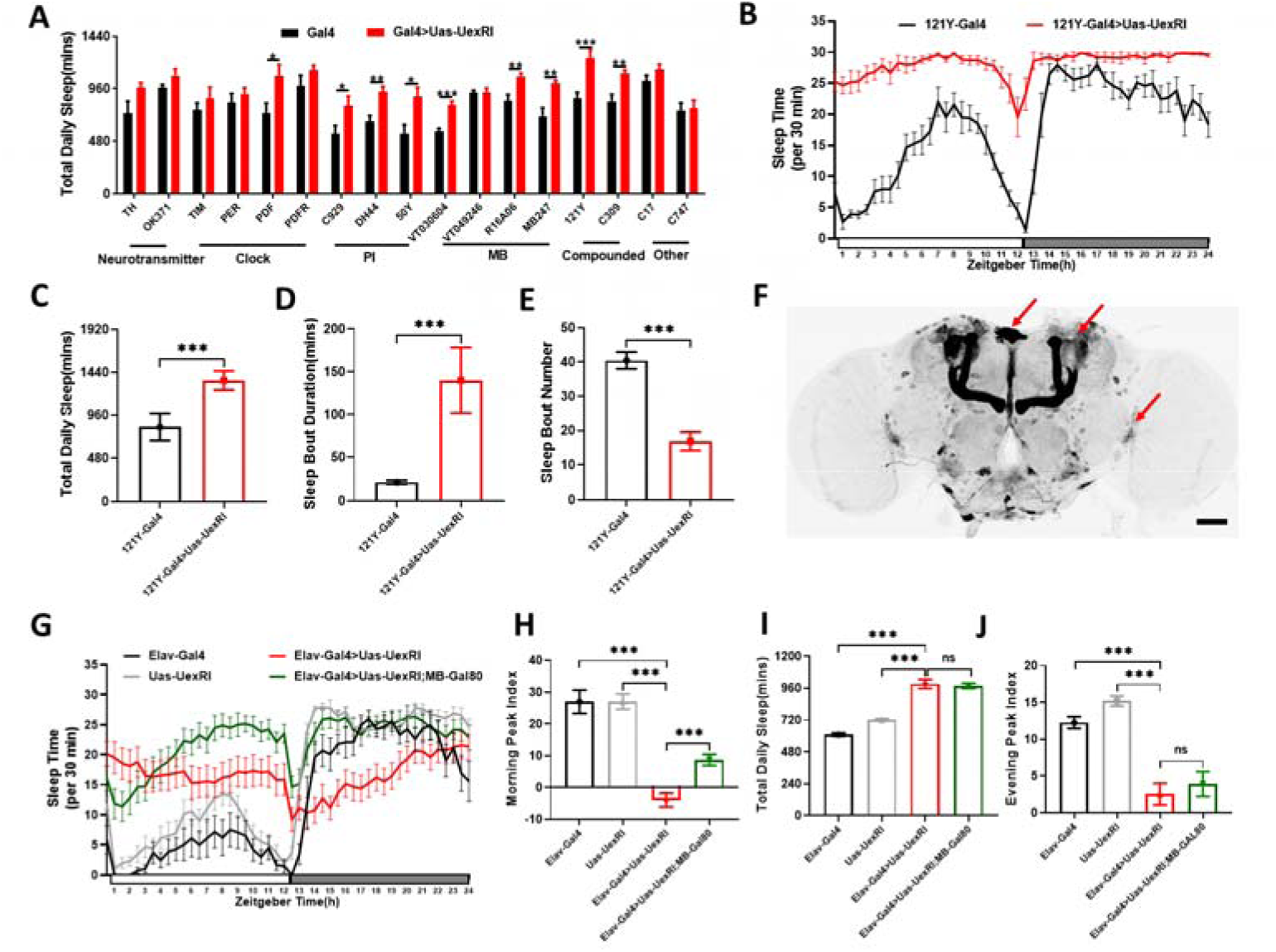
UEX functions in multiple brain regions containing PDF, MB and PI neurons. (A) The daily total sleep amount for control flies (black column, n=8) and Uex RNAi flies (red column, n=8). The Gal4 lines are grouped and labeled according to expression area. (B) Typical sleep profile of 121Y-Gal4 flies (black line, n=16) and 121Y-Gal4 > UAS-Uex RNAi flies (red line, n=16). Sleep time was plotted in 30-min bin. The x-coordinate represents zeitgeber time and the y-coordinate represents sleep time every 30 minutes. (C) The total daily sleep quantification in (B). (D) The mean sleep bout duration quantification in (B). (E) The number of quantified sleep bouts in (B). (F) Whole-mount brain immunostaining of a 121Y-Gal4 > UAS-CD8::GFP animal with anti-GFP. Arrows indicate PI, MB and PDF neurons. Scale bar = 50μm. (G) Typical sleep profile of Elav-Gal4 flies (black line, n=8), UAS-Uex RNAi flies (gray line, n=8), Elav-Gal4 > UAS-Uex RNAi flies (red line, n=13) and Elav-Gal4 > UAS-Uex RNAi; MB-gal80 flies (green line, n=13). Sleep time was plotted in 30-min bin. The x-coordinate represents zeitgeber time and the y-coordinate represents sleep time every 30 minutes. (H) The total daily sleep quantification in (G). (I) The morning peak quantification in (G). (J) The evening peak quantification in (G).

The co-immunostaining of PDF and GFP in *121Y-Gal4 > UAS-CD8::GFP* flies showed that PDF LNvs had an overlap with GFP (Figure 2–figure supplement 2A). Although Uex knockdown in PDF neurons caused sleep increase (Figure 2A, Figure 2–figure supplement 2B), we failed to notice any obvious disruption of the circadian feedback loop (Figure 2–figure supplement 2C) or circadian expressions of PDF protein (Figure 2–figure supplement 2D). We also introduced the expression of various GAL80 proteins (Suster et al., 2004) to inhibit *Elav*>*Uex* RNAi in the neurons. Among the GAL80 lines used including CHA-GAL80, VGLUT-GAL80, GAD1-GAL80, TSHGAL80and MB-GAL80, only MB-GAL80 could partially rescue the sleep phenotype (Figure 2G), with a certain recovery effect for the morning peak (Figures 2G,2H), but not for the total daily sleep time or the evening peak (Figure 2I-2J). Previous studies have reported that morning activity bouts are mainly controlled by PDF^+^ LNvs (Chatterjee and Rouyer, 2016; Grima et al., 2004). Here we found that the MB neurons, with their specific coordination of UEX protein, also play an important role in regulating morning activity. In addition, *Uex* knockdown resulted in no obvious differences in sleep recovery after sleep deprivation (Figure 2–figure supplement 2E). We infer that Uex acts in multiple brain regions containing PDF, MB and PI neurons to control total sleep time.

### 3. UEX and CNNM1 are homologous in regulating sleep

As the sleep profile of *121Y>Uex* RNAi flies was most similar to that of *Elav>Uex* RNAi flies and 121Y-GAL4 cover the expression of essential sleep promoting neurons upon Uex knockdown (Figure 1A and Figure 2B), we took advantage of the 121Y-GAL4 strain to conduct rescue experiments. *Drosophila* UEX and mammalian homologous protein CNNMs have a similarity of 70% in amino acid sequence, and greater than 90% in the conserved DUF21 and CBS domains (Figure 3–figure supplement 3). In order to verify the functional homology between UEX and CNNMs, we generated transgenic flies, and performed rescue experiments in *121Y*>*Uex* RNAi flies with overexpression of wild-type UEX, mutant UEX (T499I that is homologous to the CNNM2 T562I to cause dominant hypomagnesemia in humans (Stuiver et al., 2011)), CNNM1, CNNM2, or CNNM4. Results showed that, overexpression of wild-type UEX or CNNM1 in 121Y>*Uex* RNAi flies could rescue the sleep phenotype including the total daily sleep and the sleep bout duration, but this did not occur when overexpressing the mutated UEX T499I (Figures 3A-3C). Overexpression of CNNM2 or CNNM4 failed to rescue the phenotype and resulted in the death of the flies. Comparing the gene expression information in the open database, the tissue-specific expression of CNNM1 was seen to be significantly higher than that of other CNNM members in the human frontal cortex (Figure 3D). Similarly, the qPCR analysis of mRNA levels of CNNM1, CNNM2, CNNM3, and CNNM4 showed that only CNNM1 as highly expressed in mouse brains (Figure 3E).

**Figure 3.**
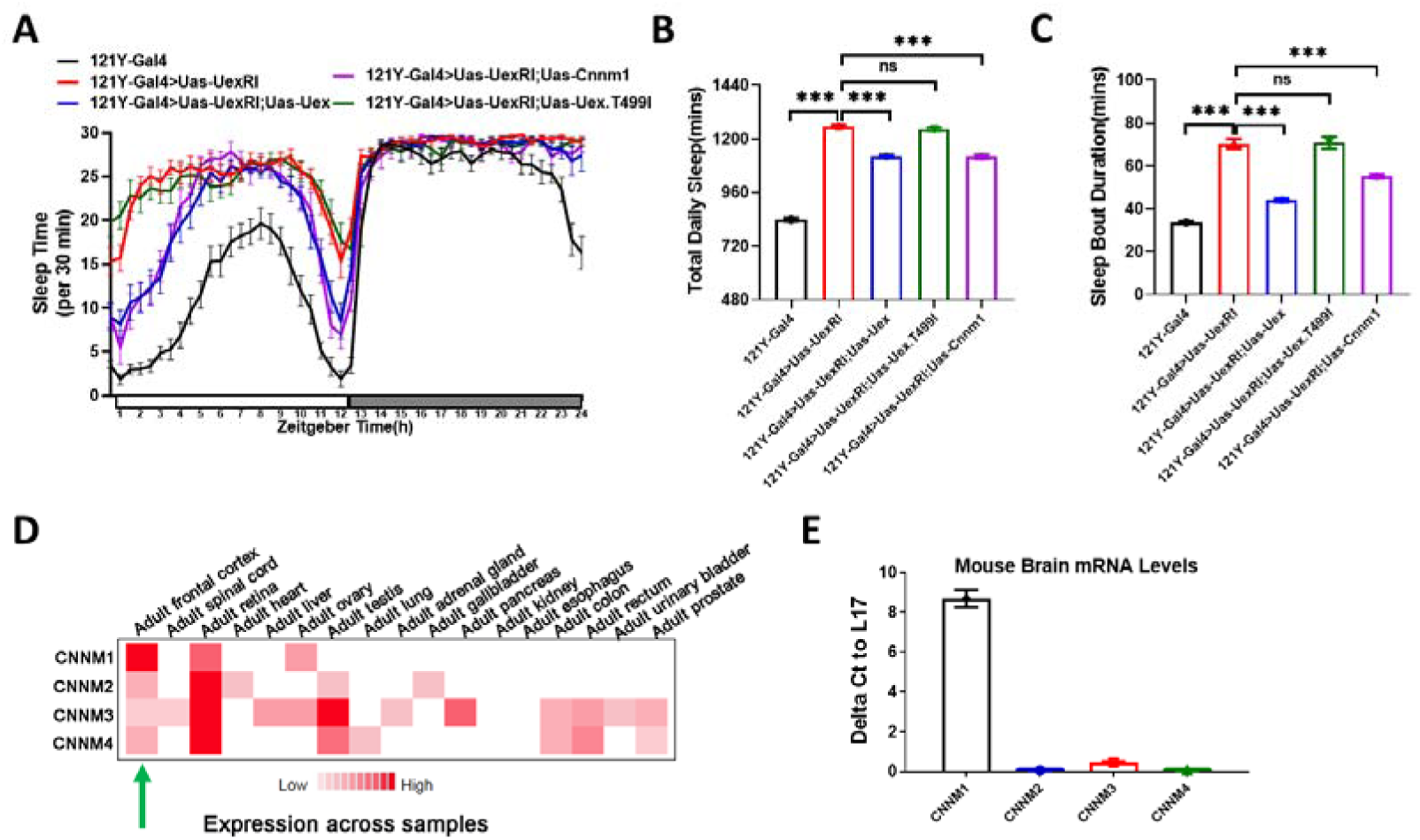
UEX and CNNM1 are homologous in regulating sleep. (A) Sleep profile of 121Y-Gal4 flies (black line, n=16), 121Y-Gal4 > UAS-Uex RNAi flies (red line, n=16), 121Y-Gal4 > UAS-Uex RNAi; UAS-Uex flies (blue line, n=16), 121Y-Gal4 > UAS-Uex RNAi; UAS-Uex.T499I flies (green line, n=16) and 121Y-Gal4 > UAS-Uex RNAi; UAS-Cnnm1 flies (purple line, n=16). Sleep time was plotted in 30-min bin. The x-coordinate represents zeitgeber time and the y-coordinate represents sleep time every 30 minutes. (B) The total daily sleep quantification in (A). (C) The mean sleep bout duration quantification in (A). (D) Tissue expression data of human CNNMs. The data derived from the Human Protein Atlas database. The human frontal cortex is indicated by the green arrow. The deeper the red color, the higher the protein expression. (E) Relative mRNA levels of cnnm1(black column, n=3), cnnm2(blue column, n=3), cnnm3 (red column, n=3) and cnnm4 (green column, n=3) to ribosomal protein L17 in mouse brain.

### 4. UEX functions as a magnesium transporter and exhibits sleep-related rhythmic oscillations

CNNMs have been reported as magnesium transporters that localize on the cell membrane (Chen et al., 2018b; Chen et al., 2020; Gulerez et al., 2016). Using *Drosophila* salivary gland cells and S2 cells to ectopic express UEX, we observed the localization of UEX-Flag on the cellular membrane (Figures 4A-4B). Considering UEX as a sleep regulator, we observed the circadian expression of UEX in the brain at the protein level, but not the mRNA level (Figures 4C-4E), which indicated that UEX may mediate circadian magnesium flux in *Drosophila* neurons. Magnesium Green (MG) is a magnesium indicator that has been developed for real-time tracking magnesium transport events (Erickson and Moerland, 2005). We utilized MG to measure the magnesium concentration of S2 cells incubated with an appropriate Mg^2+^ loading buffer (Figure 4–figure supplement 4A). Results showed that overexpression of UEX accelerated Mg^2+^ efflux as indicated by a reduction in the relative MG fluorescent intensity in the cells (Figure 4F), while overexpression of the mutant UEX-T499I had no such effect and even block the effects (Figure 4F).

**Figure 4.**
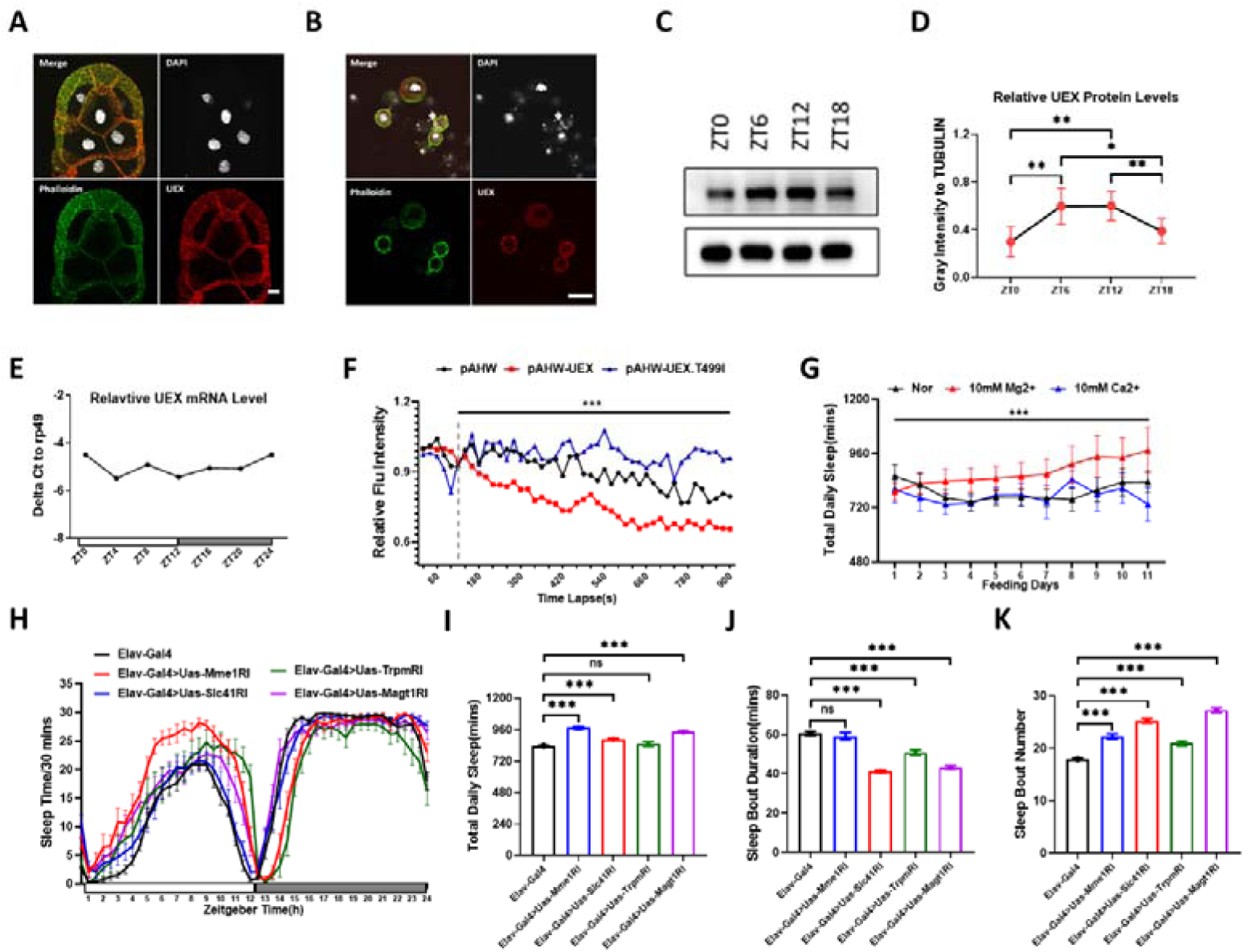
UEX functions as a magnesium transporter and exhibits sleep-related rhythmic oscillations. (A) Flag-UEX was ectopic expressed in salivary glands with 121Y-Gal4 to observe UEX localization. The nucleus was labeled with DAPI fuel (Blue), and the inner cell membrane was stained with phaneropeptide (Green). FLAG antibody was used to observe UEX localization (Red). Scale bar = 15μm. (B) Flag-UEX was ectopically expressed in Drosophila S2 cells to observe UEX localization. The nucleus was labeled with DAPI fuel (Blue), and the inner cell membrane was stained with phaneropeptide (Green). FLAG antibody was used to observe UEX localization (Red). Scale bar = 15 μm. (C) Relative protein levels of Uex in adult fly head compared to internal reference TUBULIN. Four time points were chosen to test (ZT0, Z6, ZT12, ZT18). (D) Quantification of Uex protein levels relative to Tubulin protein levels (n=3). Data are from the same flies as in Figure C. (E) Relative mRNA levels of Uex compared to rp49 in adult wild type fly heads(n=3). Six time points were chosen to test (ZT0, ZT4, ZT8, ZT12, ZT16, ZT20). The gray bar on the x-coordinate represent night time. (F) Drosophila S2 cells transfected with pAHW-3HA (black line), pAHW-3HA-UEX (red line) or pAHW-3HA-UEX.T499I (blue line) were cultured in high magnesium (40 mM) Schneider’s medium loaded with Magnesium Green (2 μM), and then the magnesium concentration is diluted to 20 mM at the indicated time point (vertical dotted line). The means of relative fluorescence intensities of 6 cells are indicated. (G) The daily total sleep amount of wild type flies fed with no iron (black line, n=13), 10 mM MgCl_2_ (red line, n=14) and 10 mM CaCl_2_ (blue line, n=8). Sleep time was plotted in 24-h bin. The x-coordinate represents day counts and the y-coordinate represents daily total sleep time. (H) Typical sleep profile of Elav-Gal4 flies (black line, n=32), Elav-Gal4 > UAS-Mme1 RNAi flies (red line, n=16), Elav-Gal4 > UAS-Slc41 RNAi flies (blue line, n=16), Elav-Gal4 > UAS-Trpm RNAi flies (green line, n=13) and Elav-Gal4 > UAS-MagT1 RNAi flies (purple line, n=18). Sleep time was plotted in 30-min bin. The x-coordinate represents zeitgeber time and the y-coordinate represents sleep time every 30 minutes. (I) The total daily sleep quantification in (H). (J) The mean sleep bout duration quantification in (H). (K) The number of quantified sleep bouts in (H).

We then wondered whether magnesium supplementation could influence sleep behavior in flies. We added magnesium at a safe concentration (10 mM) (Georgiev et al., 2010) into the food for groups of 5-day-old adult flies. Fly groups fed without magnesium were used as controls and fly groups fed with the same concentration of calcium served as a divalent cation supplemental control (Figure 4–figure supplement 4B). The sleep of all fly groups was then continuously recorded. With the increase of feeding days, flies fed continuously with magnesium supplementation showed an increase in daily total sleep, while both control groups showed normal daily sleep (Figure 4G and S4C).

To observe if any other magnesium transporters, beyond CNNMs/UEX, influence the nervous system and sleep behavior, we performed the knockdown each of the *Drosophila* homologs of the mammalian magnesium transporters MME1, SLC41, MAGT and TRPM7 using the pan-neuronal *ELAV-GAL4* driver. The knockdown of each of these magnesium transporters resulted in various effects on sleep with increases or decreases in either total daily sleep, sleep bout duration, or sleep bout number to various extents (Figures 4H-4K). Among these, knockdown of *Uex* showed a dominant effect on sleep.

### 5. UEX deficiency interferes with Ca^2+^ levels and CREB2 signaling

Mg^2+^ serves as a calcium antagonist, affecting cellular Ca^2+^ signaling (de Baaij et al., 2015; Jahnen-Dechent and Ketteler, 2012). As UEX functions to efflux intracellular magnesium we considered if *Uex* knockdown might influence calcium levels. It was interesting to note that in our experiments the supplementation of Mg^2+^ in food could not relieve or aggravate the sleep phenotype in *121Y>Uex* RNAi flies, whereas supplemental Ca^2+^ was able to partially relieve the daily total sleep of *121Y>Uex* RNAi flies (Figure 5A and S4D). Upon such evident effects of UEX knockdown on Ca^2+^ levels and neuronal activity, we applied GCaMP6s methods (Liu et al., 2016) to view the calcium levels in the calyx region of the adults fly brains at the ZT0-ZT1 and ZT12-ZT13 time points. The signals were measured and utilized the RFP expression as a comparison to exclude the possible effect of relative transcription and light emission. GCaMP6m signals were significantly reduced in the *121Y>Uex RNAi* flies during both ZT0-ZT1 and ZT12-ZT13, compared with those of controls (Figures 5B-5E).

**Figure 5.**
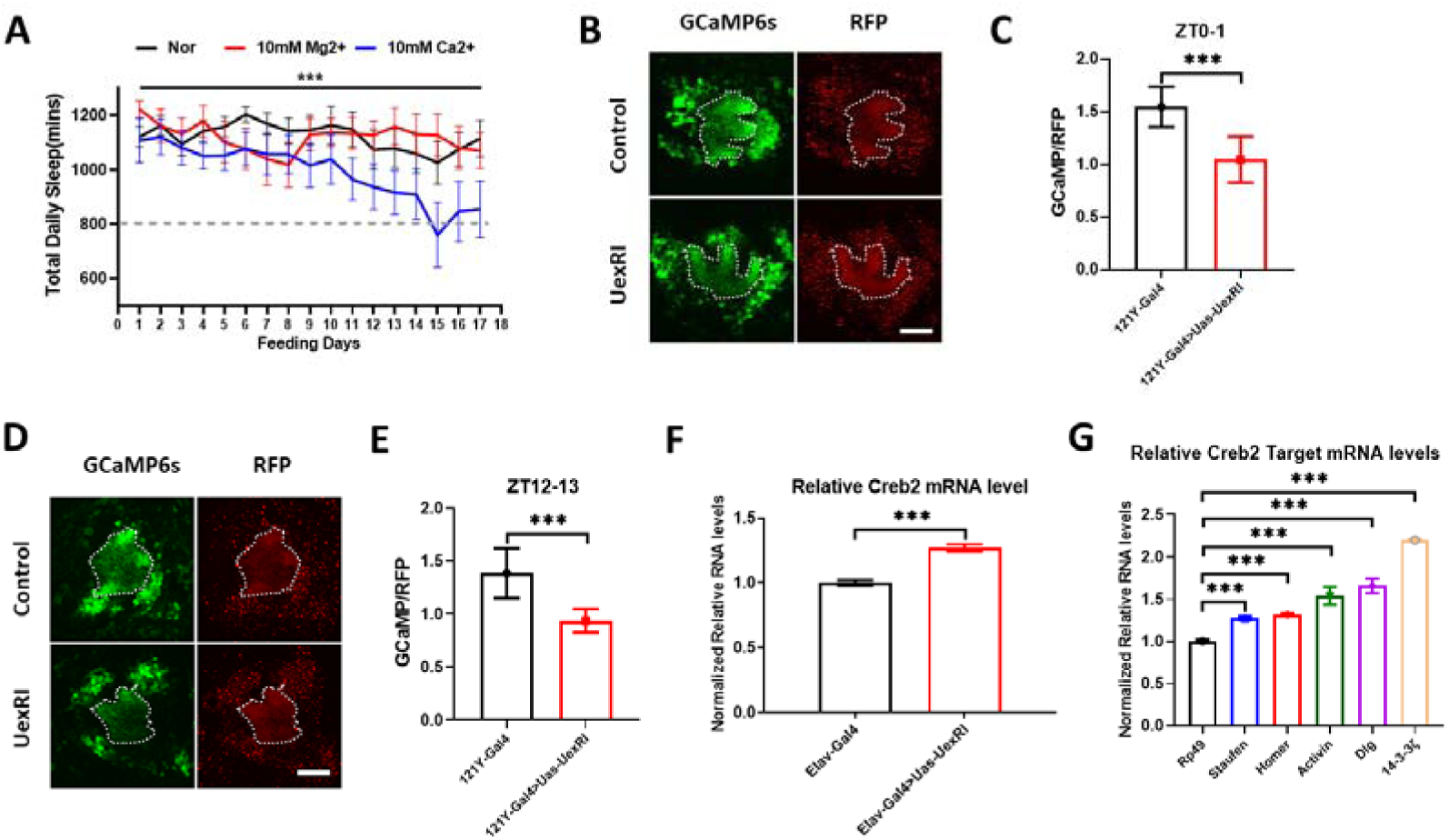
UEX deficiency interferes with Ca^2+^ levels and its related CREB2 signaling. (A) The daily total sleep amount of 121Y-Gal4 > UAS-Uex RNAi flies fed with no iron (black line, n=14), 10 mM MgCl2 (red line, n=8) and 10 mM CaCl2 (blue line, n=14). Sleep time was plotted in 24-h bin. The x-coordinate represents day counts and the y-coordinate represents daily total sleep time. The normal daily sleep time of flies was roughly plotted (horizontal gray dotted line). (B) Representative images of GCaMP (upper panels) and myrRFP (lower panels) fluorescence intensity in the nerve bundles of Calyx neurons of 121Y-Gal4 > UAS-GCaMP6m, UAS-myrRFP flies and 121Y-Gal4 > UAS-Uex RNAi, UAS-GCaMP6m, UAS-myrRFP flies at ZT0-1. Average projections of continuous 50 frames in the 3-min recording are used for quantification. Scale bar = 20μm. (C) Relative fluorescence intensity quantification of Calyx GCaMP6/myrRFP in (B). (D) Representative images of GCaMP (upper panels) and myrRFP (lower panels) fluorescence intensity in the nerve bundles of Calyx neurons of 121Y-Gal4 > UAS-GCaMP6m, UAS-myrRFP flies and 121Y-Gal4 > UAS-Uex RNAi, UAS-GCaMP6m, UAS-myrRFP flies at ZT12-13. Average projections of continuous 50 frames in the 3-min recording are used for quantification. Scale bar = 20μm. (E) Relative fluorescence intensity quantification of GCaMP6m/myrRFP in (D). (F) Relative mRNA levels of Creb2 to rp49 in Elav-Gal4 > UAS-Uex RNAi flies head (red column, n=3) compared with in Elav-Gal4 control (black column, n=3). (G) Relative mRNA levels of different neural plasticity-related genes: Staufen (blue column, n=3), Homer (red column, n=3), Activin (green column, n=3), Dlg (purple column, n=3) and 14-3-3ζ (yellow column, n=3) to control rp49 (black column, n=3) in Elav-Gal4 > UAS-Uex RNAi fly head, compared with in Elav-Gal4 control fly heads.

Uncoordinated calcium flow was seen as a catalyst for the inhibition of CREB2 transcription, thereby suppressing neuroplasticity-dependent learning and memory events in *Drosophila* (Hendricks et al., 2001; Miyashita et al., 2012; Sanyal et al., 2002). It is speculated that the disordered calcium flow caused by *Uex* knockdown may affect CREB2-dependent neuroplasticity-related gene transcription. Such targeted genes as *Staufen, Homer, Activin, Dlg*, and *14-3-3ζ* also play an important role in sleep behavior (Miyashita et al., 2012). We examined the transcription level of the *Creb2* in the head tissue of *Elav>Uex RNAi* flies which showed a significant increase, compared to the controls (p<0.01) (Figure 5F). Correspondingly, the mRNA levels of the CREB2-targeted genes were also detected to be significantly up-regulated (p<0.001) (Figure 5G).

### 6. UEX interacts with CNK to regulate sleep via CNK-dependent ERK signaling

Partially rescue effect of calcium supplement on sleep phenotype in *121Y>Uex RNAi* flies implicates a complicated role for *Uex*. We sought to search for UEX-interacting molecules that participate in the regulation of sleep time. A previous report on the ERK signaling pathway has screened out an interaction between CNK and UEX (Friedman et al., 2011). We verified protein interactions between UEX and CNK in S2 cells (Figure 6A). Using *Cnk-RNAi* lines, we also observed severe sleep increase in *Elav>Cnk RNAi* flies (Figure 6B), showing increased total daily sleep and sleep bout duration (Figure 6C-6D). Repeated experiments using another *Cnk-RNAi* line further reinforced these conclusions (Figure 6–figure supplement 5A-5D), resulting in two RNAi lines that effectively knocked down *Cnk* mRNA levels (Figure 6–figure supplement 5E).

**Figure 6.**
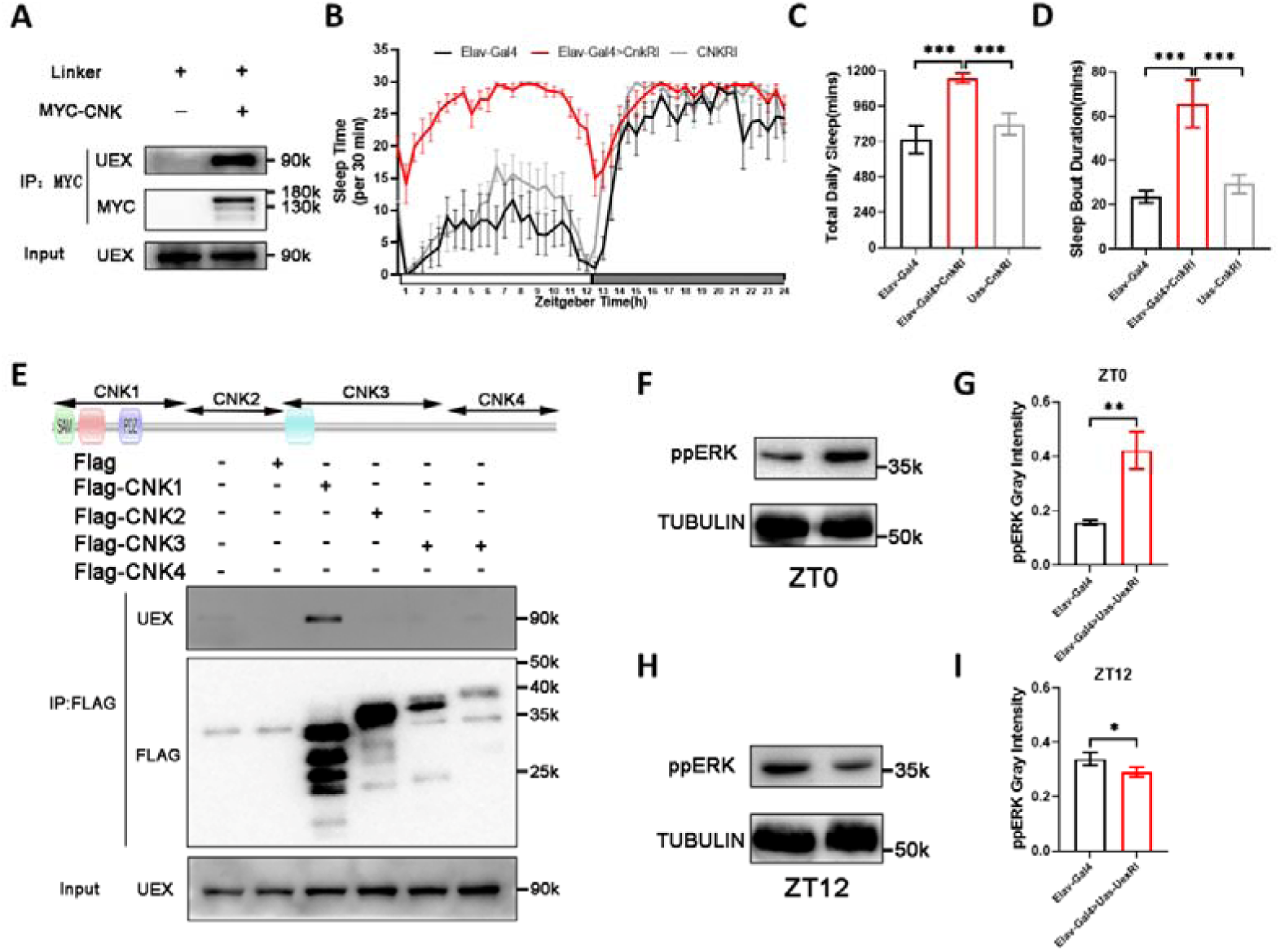
UEX interacts with CNK to regulate sleep via a CNK-dependent ERK pathway. (A) Western blot results for UEX-CNK interaction. Drosophila S2 cells were transfected with pAC5.1-MYC and pAC5.1-MYC-CNK. MYC antibodies were used to conduct the IP experiment. (B) Typical sleep profile of Elav-Gal4 flies (black line, n=16), UAS-Cnk RNAi flies (gray line, n=16) and Elav-Gal4 > UAS-Cnk RNAi flies (red line, n=16). Sleep time was plotted in 30-min bin. The x-coordinate represents zeitgeber time and the y-coordinate represents sleep time every 30 minutes. (C) The total daily sleep quantification in (B). (D) The mean sleep bout duration quantification in (B). (E) Western blot results for interaction between UEX and separate CNK domains. Drosophila S2 cells transfected with nothing, pAC5.1-FLAG, pAC5.1-FLAG-CNK1, pAC5.1-FLAG-CNK2, pAC5.1-FLAG-CNK3, or pAC5.1-FLAG-CNK4, with each part of CNK marked at the top of the graph and the different domains of CNK are indicated in the diagram. FLAG antibody was used to conduct IP experiment. (F) Relative protein levels of ppERK and TUBULIN in Elav-Gal4 fly heads and Elav-Gal4 > Uex RNAi fly heads at early morning (ZT0). (G) Quantification of ppERK protein levels relative to TUBULIN protein levels (n=3) in (F). (H) Relative protein levels of ppERK and TUBULIN in Elav-Gal4 fly heads and Elav-Gal4 > Uex RNAi fly heads at early night (ZT12). (I) Quantification of ppERK protein levels relative to TUBULIN protein levels (n=3) in (H).

The CNK protein, as a scaffold protein, interacts with S6K and RAF to suppress ERK signal transduction through the N-terminal domain, including the SAM and PDZ domains (Douziech et al., 2003). To discover which domain of CNK protein could interact with UEX, we constructed plasmids containing each of the 4 segmented domains and transfected each one into S2 cells. Co-IP results showed that among the four domains tested, only the N-terminal domain could interact with UEX (Figure 6E). ERK signaling plays an essential role in promoting sleep behavior in which the expression of phosphorylated ERK (ppERK) shows a low level in the morning (ZT0) and a high level in the evening (ZT12) in *Drosophila* brains (Foltenyi et al., 2007). We examined the protein levels of ppERK in *Uex* knockdown flies. In total contrast to that of control flies, the level of ppERK in the head tissues of *Elav*>*Uex* RNAi flies showed a high level of ppERK at ZT0 (Figure 6F-6G) and a low level of ppERK at ZT12 (Figure 6H-6I). Meanwhile, ppERK was continuously maintained at high levels.

### 7. UEX regulates sleep-related neuroplasticity and synaptic plasticity

The Calcium-dependent CREB and ERK pathways are both essential components for neuroplasticity regulation (Hendricks et al., 2001; Sanyal et al., 2002; Weislogel et al., 2013) and for coordinated sleep behavior (Mackiewicz et al., 2008). We therefore wondered whether *Uex* knockdown would affect neuroplasticity. Among *Drosophila* clock neurons, PDF neuron terminals have proved to be an excellent measure of rhythmic changes in neuroplasticity due to their easily observable morphology and obvious alterations along with the neuronal terminals that open in the morning (ZT2) and contract in the evening (ZT14) (Donlea et al., 2009; Fernández et al., 2008; Petsakou et al., 2015). Since *Uex* knockdown in PDF neuron causes a sleep increase (Figure 2A, Figure 2–figure supplement Figure 2B), we performed a specific knock-down of *Uex* in PDF neurons (*PDF>Uex RNAi*) to observe the related neuroplasticity. Control flies showed a normal phenotype with PDF neuron terminals opening in the morning and contracting at night, while *PDF>Uex RNAi* flies showed an opposite and reversed pattern of PDF neuron terminals with morning closing and evening opening (Figure 7A-7B). The neuron bifurcation calculation algorithm SHOLL was used to quantify the opening degree of the PDF neuron terminals between ZT2 and ZT14 (Fernández et al., 2008). Results showed significant changes in both neuron bifurcation degrees and the length of PDF neuronal terminals in *PDF>Uex RNAi* flies (Figure 7B-7C).

**Figure 7.**
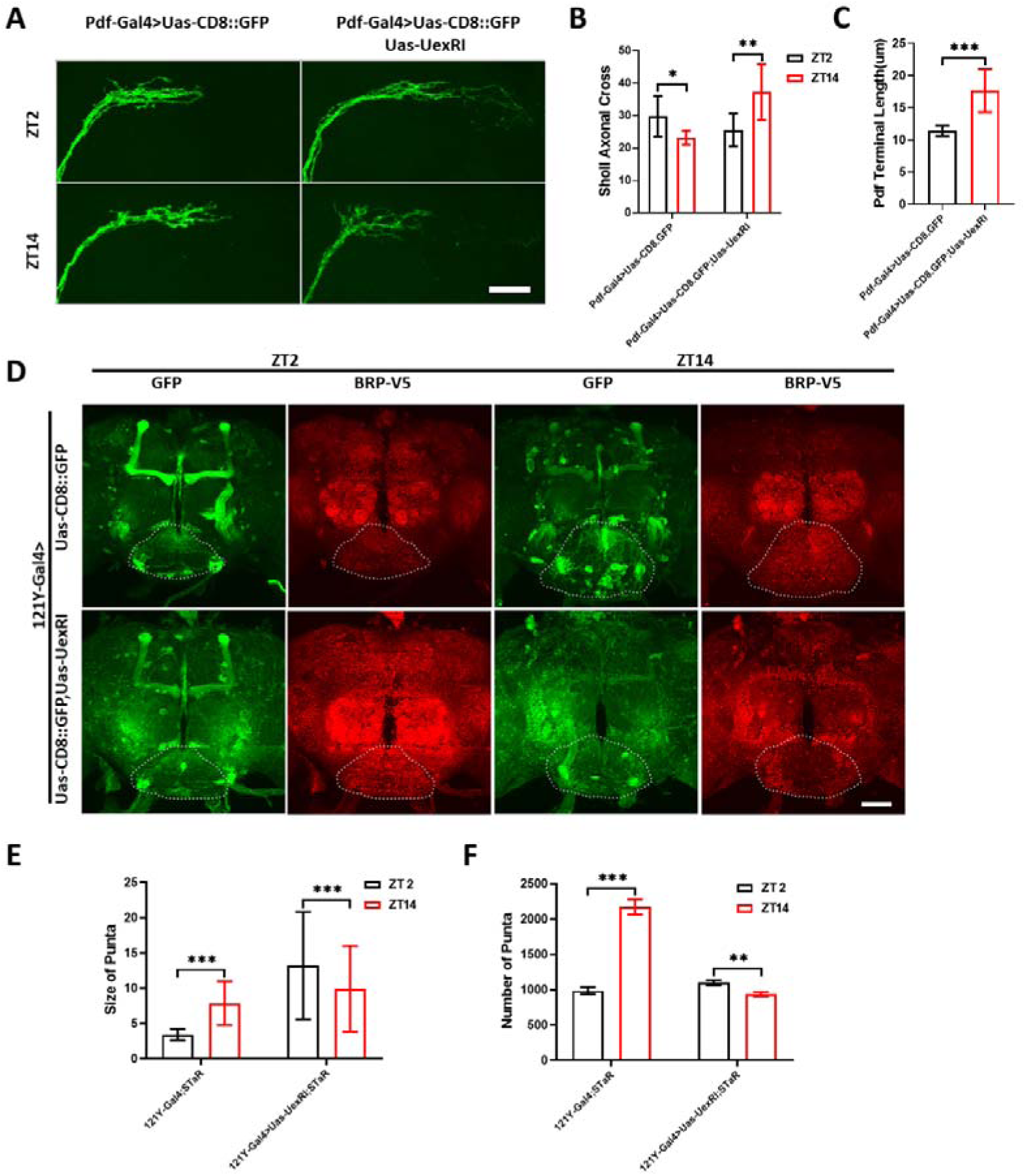
UEX regulates sleep-related neuroplasticity and synaptic plasticity. (A) Representative confocal images of nerve terminal of Pdf^+^ neurons taken during the early day (ZT2) and early night (ZT14) in Pdf-Gal4 > UAS-CD8::GFP flies and Pdf-Gal4 > UAS-Uex RNAi, UAS-CD8::GFP flies. Scale bar = 25μm. (B) Sholl analysis for total axonal projection of Pdf-Gal4 > UAS-CD8::GFP flies(n=7 at ZT2, n=7 at ZT14) and Pdf-Gal4 > UAS-Uex RNAi, UAS-CD8::GFP flies(n=9 at ZT2, n=12 at ZT14). (C) Quantification of nerve terminal from the cross center in Sholl analysis to end in (A). (D) Representative images of CD8::GFP (green) and BRP-V5(red) fluorescence intensity in the 121Y-Gal4 > Brp-stop-V5-2A-LexA,Uas-FlpD, UAS-CD8::GFP flies and 121Y-Gal4 > UAS-Uex RNAi, Brp-stop-V5-2A-LexA,Uas-FlpD, UAS-CD8::GFP flies at early morning(ZT2) and early night(ZT14). The white dotted line position is used for subsequent quantitative analysis. Scale bar = 25μm. (E) Relative size quantification of synapse in (D). (F) Relative number quantification of synapse in (D).

As a marker of neuroplasticity, synapse changes occur periodically in the brain (Bushey et al., 2011). Synaptic tagging with recombination (STaR) technology has been developed to quantify the pre-synaptic T-bar structure in the specific GAL4-expressing neurons by viewing the V5 labeled BRP protein (Chen et al., 2014). We compared the BRP-V5 signals at the PI output SEZ region in the brains of control flies and *121Y>Uex* RNAi flies at ZT2 and ZT 14 (Figure 7D) and found that the alteration patterns for both the number of synapses and the size of synaptic structures at ZT2 and ZT14 were opposite to those of the controls (Figure 7D-7F). These observations were consistent with the results of the morning and evening trends of PDF neuron terminals. Taken together, UEX may function in parallel with calcium-dependent CREB2 signaling and a CNK-ERK pathway to regulate neuroplasticity in circadian output neurons, thus explaining why UEX loss results in neural disorders and increased sleep time.

## Discussion

The relationship between magnesium ions and their transporter proteins, and in their involvements in regulating specific neuronal behaviors such as sleep, have been poorly reported. More generally, magnesium has certainly been recognized for its important pharmacological roles. Magnesium sulfate, for example, is used as a second line drug for migraine patients (Corbo et al., 2001), to aid sleep in cases of depression (Serefko et al., 2016), or to relax nerves or muscle tension in cases of preeclampsia (Duley et al., 2010). Here we report that the magnesium transporter UEX/CNNM plays an important role in regulating sleep. Disturbance of the rhythmically expressed UEX in the nervous system caused a sleep disorder. The mechanisms include that UEX functions both in maintenance of neuroplasticity by regulating the Mg^2+^/Ca^2+^ mediated CREB pathway and via ERK signal transduction by physically interacting with CNK (Figure 8).

**Figure 8.**
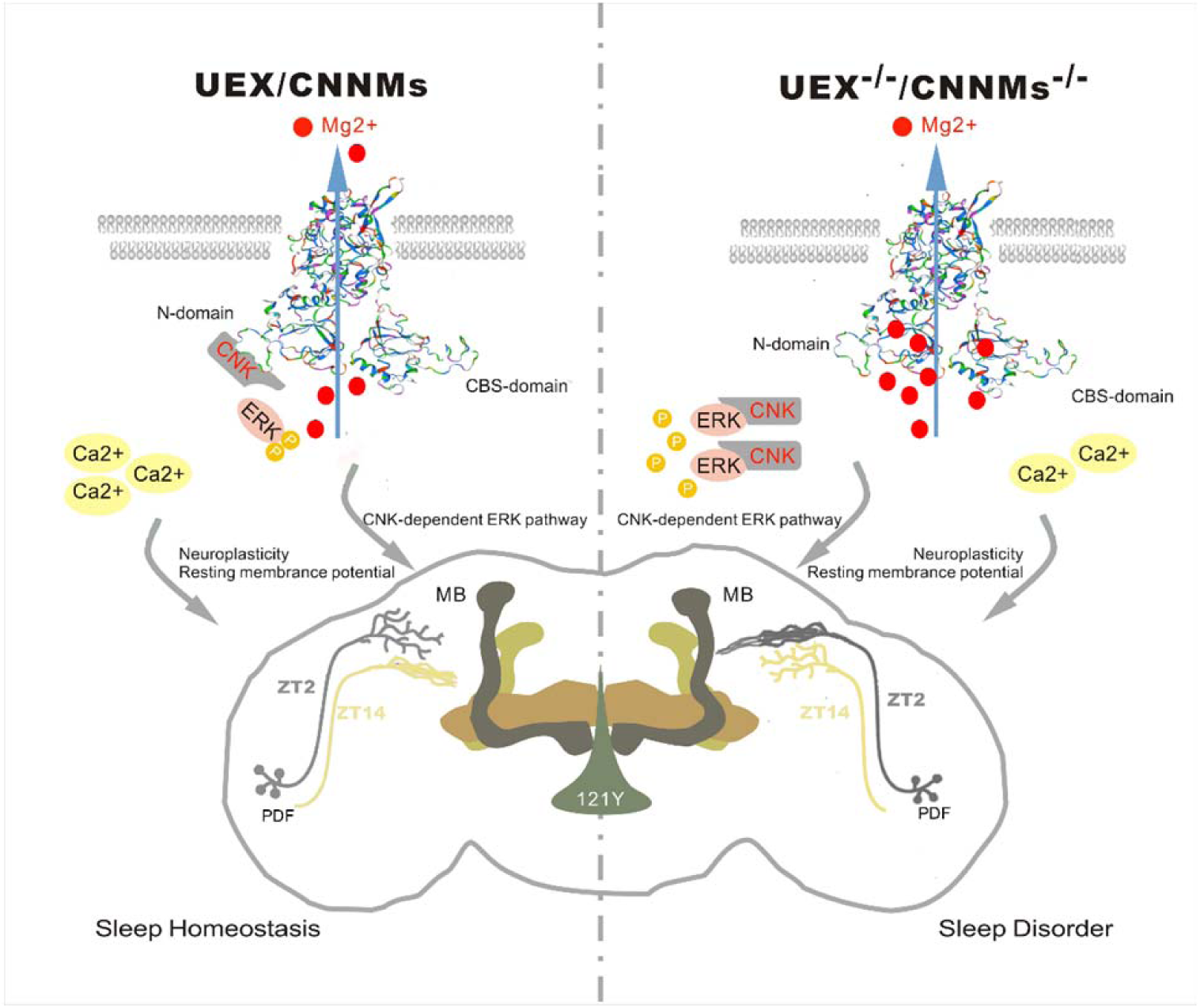
Graphic illustration. The magnesium transporter UEX regulates sleep by affecting Mg^2+^/Ca^2+^-CREB signaling and CNK-ERK pathway, and neuroplasticity. Knockdown of Uex in the nervous system results in a severe sleep disorder.

Most of the magnesium ions in the cell are in a fixed and bound state, with few free magnesium ions present. This distinguishes the activity of magnesium and calcium ions (Jahnen-Dechent and Ketteler, 2012). Whilst existing research has mainly focused on the regulation of behavior via the role of sodium and potassium in the maintenance of neuronal action potential (Cirelli et al., 2005; Kempf et al., 2019), our study suggests that magnesium also plays roles of comparable importance to sodium and potassium in neural circuits. However, magnesium does not display obvious differences between intracellular and external concentrations (de Baaij et al., 2015), a situation also highly differing from that of either sodium or potassium ions. This suggests that effects of magnesium ions on neurons are probably not executed via rapid modifications of membrane potential, but facilitated by changing the baseline value of membrane potential and intracellular calcium responses.

CNNM has been shown to influence intracellular calcium response in spermatogenesis (Yamazaki et al., 2016). Whether such an effect is direct or indirect remains unknown. We have shown that the effect of UEX on neurons is indirect, where calcium feeding does not produce any phenotype in wild-type animals, but can rescue the sleep phenotype of *Uex* knockdown flies. UEX participates in signal transduction by affecting magnesium ions to operate calcium flux. We found that alongside UEX-mediated neuroplasticity through a Ca^2+^ dependent CREB pathway, UEX also physically interacts with CNK to affect the ERK-dependent neuroplasticity pathway. Flies with *Cnk* knockdown also showed increased sleep time. The ERK signaling pathway is the authoritative signal molecule that affects neuroplasticity. Our work also reveals that UEX affects neuroplasticity at a molecular level.

A recent study found that PRL-1 knockdown in PDF neurons dramatically lengthens the circadian period in *Drosophila* (Kula-Eversole et al., 2021). UEX interacts with PRL-1(Guo et al., 2019). Here we show that *Uex* knockdown also led to significant structural alterations in PDF neuron terminals and in synaptic plasticity in *Drosophila* brains. It is possible that knockdown of PRL-1 induced a circadian phenotype that might be mediated by *Uex*. Future study may need to consider the combined functions of both UEX and PRL-1 in relating sleep homeostasis control.

In conclusion, this study demonstrates magnesium transporter UEX exhibits rhythmic oscillations in the brain. Knockdown of Uex not only affects the efflux of intracellular Mg^2+^, but also Ca^2+^-related CREB signaling and neuroplasticity, resulting in a sleep disorder. We also found that the UEX interacts with CNK and affects the circadian expression of ppERK. Knock down of CNK also results in a similar sleep phenotype. Ectopic-expression of human homolog CNNM1 can rescue the sleep phenotype in *Uex* knockdown flies, suggesting that human magnesium transporter may play a similar role in sleep regulation.

## Methods

### Experiment Conditions and *Drosophila* Stocks

Flies were raised in standard fly medium consisting of yeast, corn, and agar and at 25 °C. Virgin female flies were collected shortly after eclosion and kept at no more than 20 flies per vial to exclude social enrichment. All flies, except special lines, were kept on a 12-12 day-night cycle before experiments. 4-10-day old non-mated female flies were used for experiments.

The following stocks were obtained from the Bloomington *Drosophila* Stock Center (BDSC): C929-Gal4 (BS25373); 121Y-Gal4 (BS30815); 50Y-Gal4 (BS30820); Pdfr-Gal4 (BS33070); C17-Gal4 (BS39690); R16A06-Gal4 (BS48706); C747-Gal4 (BS6494); C309-Gal4 (BS6906); Tim-Gal4 (BS7126); Per-Gal4 (BS7127); UAS-Flp.d, brp (FRT.Stop) V5-2A-LexA-VP16/TM6B (BS55751) and MB-Gal80 (BS64306).

The following stocks were obtained from the Vienna Drosophila Research Center (VGRC): VT030604-Gal4 (v200228) and VT049246-Gal4 (v205379).

The following stocks were obtained from the Tsinghua University *Drosophila* Center: Elav-Gal4 (TB00041); UAS-Trpm RNAi (TH03911.N); UAS-Mmei RNAi (TH03984.N); UAS-Magt1 RNAi (THU1322); UAS-Slc41A RNAi (THU3850); UAS-Cnk RNAi (THU0733); UAS-Cnk RNAi (THU4863); UAS-RFP (THJ0109); and UAS-CD8::GFP (THJ0080).

PDF-Gal4 and UAS-GCaMP6m lines were obtained from Zhefeng Gong Lab in Zhejiang University. DH44-Gal4 came from Liming Wang Lab in Zhejiang University.

The following lines were from this paper or from the stocks of our own laboratory: Elav-GeneSwitch-Gal4; MB247-Gal4; OK371-Gal4; TH-Gal4; UAS-Uex; UAS-Uex.Flag; UAS-Uex.T499I; UAS-CNNM1.

### Sleep Assays and Related Data Analysis

*Drosophila* sleep, as monitored using the *Drosophila* Activity Monitoring System (DAM) and analyzed using the MATLAB program (Donelson et al., 2012), is defined as the state where the number of activities is zero within five minutes. Briefly, flies were collected into polycarbonate tubes with 2% (m/V) agar medium food and assembled into the DAM system. For sleep deprivation experiments, flies were shaken violently for 10-sec at every three-minute interval, with the shaking period lasting for 12 hours. After 12h deprivation, sleep was monitored for 24 hours. All recording procedures and data collection were performed under 12-h light: 12-h dark cycle. Data were collected with a 1-min bin and summarized to 30-min bins for analysis. Statistical analysis was conducted using the Vecsey Sleep and Circadian Analysis MATLAB Program (SCAMP) 2019_v2.

### RNA isolation and Real-time Quantitative PCR

Total RNA was isolated using Trizol reagent (Thermo Fisher Scientific) from adult fly heads. cDNA was synthesized from RNA samples by using a First-Strand cDNA synthesis kit (Thermo Fisher Scientific). Real-time PCR was performed on an ABI7900HT fast real-time PCR machine. The primer for Activin, Homer, Staufen, Dlg, 14-3-3ζ and dCREB2 was same as other study (Miyashita et al., 2012). The following PCR primers were used:

Rp49: forward, 5’-GAATTATGCATTAGTGGGA-3’;

reverse, 5’-GAATCCGGTGGGCAGCATGT-3’;

Uex: forward, 5’-GGATTTTCAGCACTTTACTC-3’;

reverse, 5’-CCATTGACATTAAACCCAAG-3’;

Timelss: forward, 5’-GGCAGTGGCGATTCCAGCCC-3’;

reverse, 5’-GTGTCTCTGGTCCTCGTAAT-3’;

Cycle: forward, 5’-TTCGCAACTCCACAGTAC-3’;

reverse, 5’-AGGGATTCTTGAAGGCC-3’;

Clock: forward, 5’-GTCAGTTCGCAAAGCCA-3’;

reverse, 5’-CGGCTCAAGAAATGTCG-3’;

L17: forward, 5’-CGGTATAATGGTGGAGTTG-3’;

reverse, 5’-ACCCTTAAGTTCAGCGTTACT-3’.

Cnnm1: forward, 5’-TTGTCCAAAGAGCCGTCCTG-3’;

reverse, 5’-CTAGGATGCTTTGGAAGTGC-3’.

Cnnm2: forward, 5’-TTGTCCAAAGAGCCGTCCTG-3’;

reverse, 5’-CTAGGATGCTTTGGAAGTGC-3’.

Cnnm3: forward, 5’-TTGTCCAAAGAGCCGTCCTG-3’;

reverse, 5’-CTAGGATGCTTTGGAAGTGC-3’.

Cnnm4: forward, 5’-TTGTCCAAAGAGCCGTCCTG-3’;

reverse, 5’-CTAGGATGCTTTGGAAGTGC-3’.

Cnksr1: forward, 5’-GGGTTAACTAGGGAGCTGGC-3’;

reverse, 5’-AGATTGATAAGCCCTTCGGC-3’.

Cnksr2: forward, 5’-CAGGTTGTGGCATAGGAGGA-3’;

reverse, 5’-TGCTGCAAACAGTCATCAAGA-3’.

Cnksr3: forward, 5’-ATGAGAATGTGAGTTTCGGCTA-3’;

reverse, 5’-AATGGTAAATCTGCGTCTTTGG-3’.

Cnk: forward, 5’-GAAACTTCCATTATCACCTG-3’;

reverse, 5’-GCATTTGAGCACATTGCCAC-3’.

Activin: forward, 5’-TGGCAAAAATGGTGAGATGA-3’;

reverse, 5’-TCCAATGCTAGAGCGACCTT-3’.

Homer: forward, 5’-CGAACAACCGATTTTCACCT-3’;

reverse, 5’-CTAACGTTGACCGCCTTCAT-3’.

Staufen: forward, 5’-CACCAACCAACGAAACACAG-3’;

reverse, 5’-GTTGCTACCATGGGCACTTT-3’.

Dlg: forward, 5’-CCACCACAACTTGGACACTG-3’;

reverse, 5’-ATTTGTTGCTGCTGCTGTTG-3’.

14-3-3ζ: forward, 5’-AAAACCAAAATGCAGCCAAC-3’;

reverse, 5’-CCAATTGGCAAGCTTTGTCT-3’.

Creb2: forward, 5’-CCAATGACGTGGTCGATGT-3’;

reverse, 5’-CTTTGTGGGTTCTGTTGCTG-3’.

For the quantification, the control data was normalized to average as 1 based on the Ct of control genes and the experimental data was adjusted based on the change of control data. 3 repeats were summarized into group for data presentation.

### S2 cell culture

*Drosophila* S2 cells were cultured in Schneider’s medium (Sigma, S0146) containing 10%-20% fetal bovine serum (Gemini, 900-108) at 25°C. Plasmids were transfected with X-tremeGene HP (Roche Applied Science) for imaging, immune staining, Western blot and Co-IP.

### Mg^2+^imaging analyses with Magnesium Green

Mg^2+^ imaging analyses with Magnesium Green was based on mammalian MG experiments(Yamazaki et al., 2013) and performed as follows. Before imaging, *Drosophila* S2 cells were incubated with a high Mg^2+^ loading buffer (78.1mM NaCl, 5.4mM KCl, 1.8mM CaCl_2_, 40mM MgCl_2_, 5.5mM glucose, 5.5mM HEPES-KOH, pH=7.4), including 2 mM Magnesium Green-AM (Invitrogen, M3735), for 45 min at 25°C. The cells were observed under an inverted Olympus IX81-FV1000 fluorescence microscope (Core facility, Zhejiang University School of Medicine). For continuous imaging, fluorescence of MG was measured every 20 sec. At the sixth cycle of 20 sec, the cells were diluted with dilute buffer (MgCl_2_ in the loading buffer was replaced with 60mM NaCl) to 20mM Magnesium in extracellular environment. Each recording lasted for 900 sec. Data was analyzed using Fiji/ImageJ software and presented as line plots (mean of 6 cells). The speed of magnesium transport was represented by relative fluorescence intensities after background subtraction.

### Western blot and Co-immunoprecipitation

Western blot (WB) analysis was based on the previous description (Yuan et al., 2019). Adult fly heads were pinched off and homogenized in a RIPA lysis buffer (50mM Tris-HCl, pH=8.0, 150mM NaCl, 1% (v/v) SDS, 0.5% (w/v) sodium deoxycholate, complete protease inhibitor cocktail, and PhosStop phosphatase inhibitor cocktail) at the required time point. After high-speed centrifugation the supernatant was collected, and SDS-PAGE was carried out according to the standard protocol.

For Immuno-Precipitation, S2 cells were lysed in modified RIPA lysis buffer (50mM Tris-HCl, pH=8.0, 150mM NaCl, 1% (v/v) NP40, 0.5% (w/v) sodium deoxycholate, complete protease inhibitor cocktail, and PhosStop phosphatase inhibitor cocktail). MYC-tagged or FLAG-tagged recombinant proteins were captured using ~5μg antibody and Protein-Agarose beads (Roche Applied Science, 11243233001) according to the manufacturer’s protocol. After 3-time washes with lysis buffer, SDS-PAGE was carried out according to the standard protocol.

The quantitation of the WB gray scale was completed by Fiji/ImageJ, and the ratio of target protein gray scale to TUBULIN gray scale after background subtraction was finally presented.

### Immunofluorescence and antibodies

Immunofluorescence experiments were conducted as previously described (Yuan et al., 2019). Generally, adult fly heads were dissected in cold PBS (137mM NaCl, 2.7mM KCl, 10mM Na_2_HPO_4_, 2mM KH_2_PO_4_, pH=7.4) and processed for standard wholemount immunostaining. The following antibodies were used: Mouse anti-PDF (1:200, Developmental Studies Hybridoma Bank C7); Mouse anti-Bruchpilot (1:200, Developmental Studies Hybridoma Bank nc82); Chicken anti-GFP (1:500, Abcam ab13970); Mouse anti-V5 (1:500, ThermoFisher R960-25); and Rabbit anti-FLAG (1:500, Sigma-Aldrich F2555). Phalloidin 488 (1:1000, Invitrogen A12380) was used to stain for F-actin. DAPI (1 g/ml; Sigma-Aldrich, St. Louis, MO, USA) was used to stain for nuclei.

All commercial secondary antibodies used were from the Jackson Immuno Research Laboratories. Images were obtained using a Zeiss LSM 510 upright and Olympus FV1000 confocal microscope and were processed using Adobe Photoshop (version CS6), PowerPoint 2019 Professional and Fiji/ImageJ.

### Measurements of GCaMP6m signal

121Y-Gal4> UAS-GCaMP6m; UAS-myrRFP and 121Y-Gal4> UAS-UEX RNAi; UAS-GCaMP6m; UAS-myrRFP were used for measuring intracellular Ca^2+^ levels. The axonal bundles of calyx neurons were selected as the measuring region due to consideration of signal strength and degree of signal overlap. The insensitive myrRFP signal was used to balance the basal transcription and the basal signal strength. Persistent GCaMP6m signals (50 records of single layer in 3 min) at different time points (ZT0-1 and ZT12-13) were averaged to measure the average strength of the intracellular Ca^2+^ level (Liu et al., 2016).

For experimental detail, all brains were quickly dissected in adult hemolymph-like saline (AHLS) and imaged using a Zeiss LSM 510 upright and Olympus FV1000 confocal microscope. The calyx structure was located at less than 30 sec by fast pre-scanning of the RFP signal. The 512 x 512 pixels single sections containing the nerve bundle of Calyx cells with roughly same size were selected. The ROIs of the nerve bundle were outlined by using the RFP signal. Confocal images were split by channel and the background was auto-subtracted in Fiji/ImageJ. The fluorescent signal of GCaMP6m and RFP in the same ROI were calculated using a Time Series Analyzer. Data was imported into GraphPad Prism for average processing and the results were calculated as the ratio of GCaMP6m signal to RFP signal.

### Quantitation of the bifurcations in PDF neurons

PDF-Gal4>UAS-CD8::GFP and PDF-Gal4>UAS-Uex RNAi; UAS-CD8::GFP were used to label PDF^+^ neurons and axonal branches were acquired under 400x magnification and at 800 x 800 pixels. ZT2 and ZT14 were chosen as the best times to measure the effects of sleep and wakefulness on synaptic structure. Adaptive Sholl analysis in Fiji/ImageJ was used to quantify the axonal arbor (Fernández et al., 2008). The first distinct branch was defined as the center of the circles with concentric circles spaced at 10μm. The total number of crossings between the GFP signal and all concentric circles were counted.

### BRP Measurements

121Y-Gal4>STaR; UAS-myrRFP and 121Y-Gal4>UAS-Uex RNAi; STaR; UAS-myrRFP were used for BRP measurements. ZT2 and ZT14 were chosen as the best times to measure the effects of sleep and wakefulness on synaptic structure (Fernández et al., 2008). V5 and GFP antibodys were used for immunostaining. Images were taken under 400x magnification and at 800 x 800 pixels, and further analyzed in Fiji/ImageJ. The superposed pixels of a stack and the comprising regions of interest (ROIs) were calculated. The background was subtracted before calculation and the threshold was manually determined for each stack, the standard of which was selected to keep the structure consistent. The number and size of BRP puncta were automatically calculated using the “3D Objects Counter” Plugin (Chen et al., 2014; Liu et al., 2016).

## Statistical analysis

The sleep data was auto-calculated by MATLAB software that error bar means Standard Error of Mean (S.E.M.) including the subsequent quantification of morning and evening peak. The error bar in qPCR experiments also means S.E.M.. The error bar in WB quantification, GCaMP quantification, Pdf axon quantification and Brp quantification means Standard Deviation (S.D.).

The statistical differences were calculated by analysis function of GraphPad Prism 9.0.0 software. For simple comparation between control and experimental group, the experimental group was individually compared to the control with unpaired Student’s t-test with Welch’s correction due to Gaussian data distribution. For comparison of continuous time-series data between two groups in sleep deprivation experiments, experimental data at each time point was compared to control with unpaired Student’s t-test with Welch’s correction due to Gaussian data distribution and then summarized. For comparison of continuous time-series data between three groups in magnesium green imaging and iron feeding sleep quantification, data was compared with unpaired one-way ANOVA with Welch’s correction due to Gaussian data distribution.

For all figures “*”, “**”, “***” and “****” indicate p<0.05, 0.01, 0.001 and 0.0001, respectively, and “ns” means “not significant”.

## Acknowledgement

We thank Gero Miesenboeck of Oxford University for discussions. We thank Liming Wang, Fang Guo, Wanzhong Ge, Zhefeng Gong and Fan Yang for their help on experimental equipment and techniques, Pengfei Guo, Yi Rao, Yan Zhu and Yong Ping and the Bloomington *Drosophila* Stock Center, the Exelixis Collection at the Harvard Medical School and the Tsinghua Fly Center for fly stocks. This study was supported by grants from National Key R&D Program of China (2013CB945601).

## Ethics

Animal experimentation: All of the animals were handled according to approved protocol (ZJU20160019) of the Animal Experimentation Ethics Committee of Zhejiang University.

## Data Availability Statement

All data contained in this study is available as Source Data. Source data files have been included for Figure 1, Figure 2, Figure 3, Figure 4, Figure 5, Figure 6, Figure 7, Figure 1-figure supplement 1, Figure 6-figure supplement 5.

## Author Contributions

XY, HZ, XX and HD performed the experiments, YX and XhY sourced funding and contributed to experimental design. XY and YX interpreted the data and drafted the manuscript. All authors contributed to final manuscript editing and approved the final version of the manuscript.

## Figure Supplements

**Figure 1–figure supplement 1.**
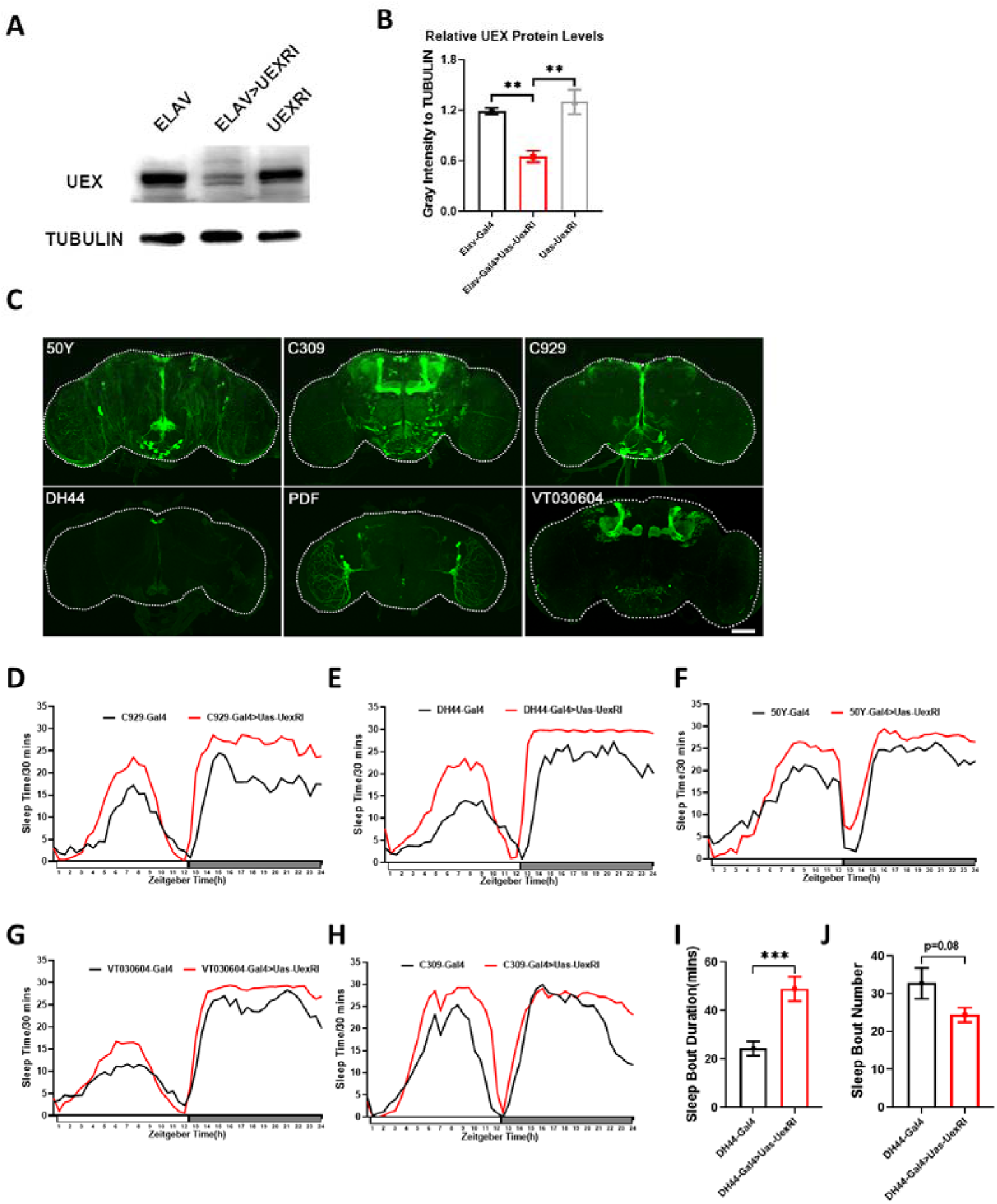
Uex function in different neurons. (A) Western Blot assay was performed to detect Uex protein expression. The TUBULIN was used as an internal reference. (B) Quantification of Uex protein levels relative to TUBULIN protein levels (n=3) in (A). (C) The pattern of drivers whose expression of Uex RNAi significantly lead to total sleep time increase in Figure 2A. 50Y-Gal4 >Uas-CD8::GFP, C929-Gal4 >Uas-CD8::GFP, DH44-Gal4 >Uas-CD8::GFP showed clear expression in PI. C309-Gal4 >Uas-CD8::GFP and VT030604-Gal4 >Uas-CD8::GFP showed clear expression in MB. Pdf-Gal4 >Uas-CD8::GFP is expressed in circadian ventral later neurons. Scale bar = 50μm. (D) Typical sleep profile of C929-Gal4 flies (black line, n=8) and C929-Gal4 > UAS-Uex RNAi flies (red line, n=8). Sleep time was plotted in 30-min bin. The x-coordinate represents zeitgeber time and the y-coordinate represents sleep time every 30 minutes. (E) Typical sleep profile of DH44-Gal4 flies (black line, n=8) and DH44-Gal4 > UAS-Uex RNAi flies (red line, n=8). (F) Typical sleep profile of 50Y-Gal4 flies (black line, n=8) and 50Y-Gal4 > UAS-Uex RNAi flies (red line, n=8). (G) Typical sleep profile of VT030604-Gal4 flies (black line, n=8) and VT030604-Gal4 > UAS-Uex RNAi flies (red line, n=8). (H) Typical sleep profile of C309-Gal4 flies (black line, n=8) and C309-Gal4 > UAS-Uex RNAi flies (red line, n=8). (I) The mean sleep bout duration quantification in (E). (J) The number of quantified sleep bouts in (E).

**Figure 2–figure supplement 2.**
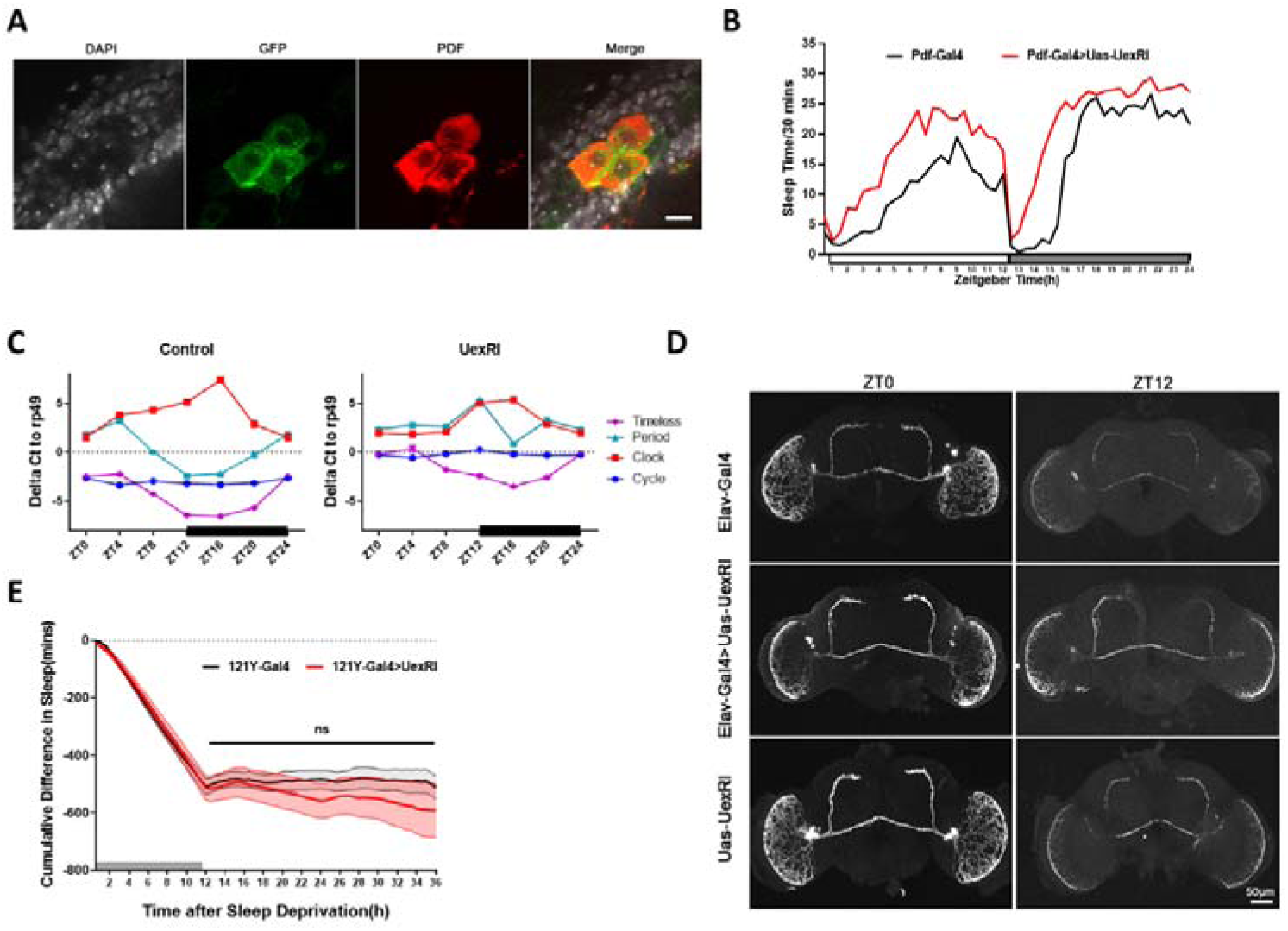
Uex functions in different neurons. (A) Immunofluorescence co-staining of GFP and PDF signal in 121Y-Gal4 > UAS-CD8::GFP. PDF positive LNv were magnified and overlap between PDF signal and GFP signal was observed. DAPI was used to stain DNA. Scale bar = 15μm. (B) Typical sleep profile of Pdf-Gal4 flies (black line, n=8) and Pdf-Gal4 > UAS-Uex RNAi flies (red line, n=8). (C) Relative mRNA levels of circadian genes to rp49 in Elav-Gal4 > UAS-Uex RNAi flies head (red column, n=3) compared with in Elav-Gal4 control (black column, n=3) at different time points (ZT0, ZT4, ZT8, ZT12, ZT16, ZT20). (D) Representative PDF protein staining images of ELAV-GAL4 control flies (upper), ELAV>Uex RNAi flies(middle) and UAS-Uex RNAi control flies(lower) at early morning (ZT0) and early evening (ZT12). Scale bar = 50 μm. (E) Continuous sleep loss quantification for 121Y-GAL4 control flies (black lines, n=14) and 121Y>Uex RNAi flies (red lines, n=13) in sleep deprivation recovery experiment. Sleep time was plotted in 30-min bin. Sleep deprivation lasted 12 hours (gray bar). The x-coordinate represents monitoring number and the y-coordinate represents accumulated sleep time loss.

**Figure 3–figure supplement 3.**
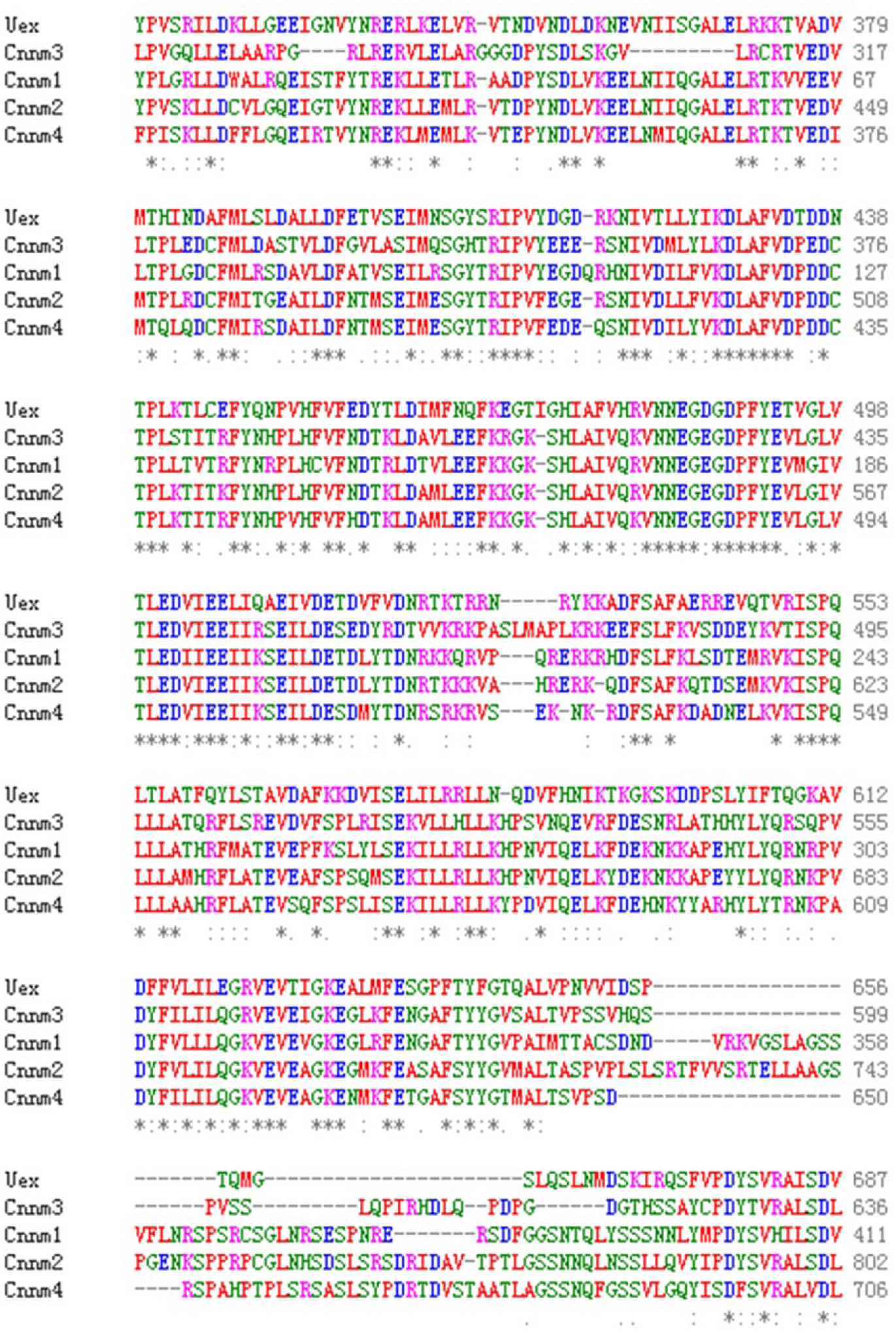
Comparison of functional homology between Uex and CNNMs families. Main amino acid alignment between UEX and CNNMs. The comparison is done through the EBI-Kalign program. “*” denotes that the sequence is consistent; “: ” denotes that the sequence is extremely conservative; “. ” denotes that the sequence is relatively conservative. The corresponding missing sequence is replaced by “-”.

**Figure 4–figure supplement 4.**
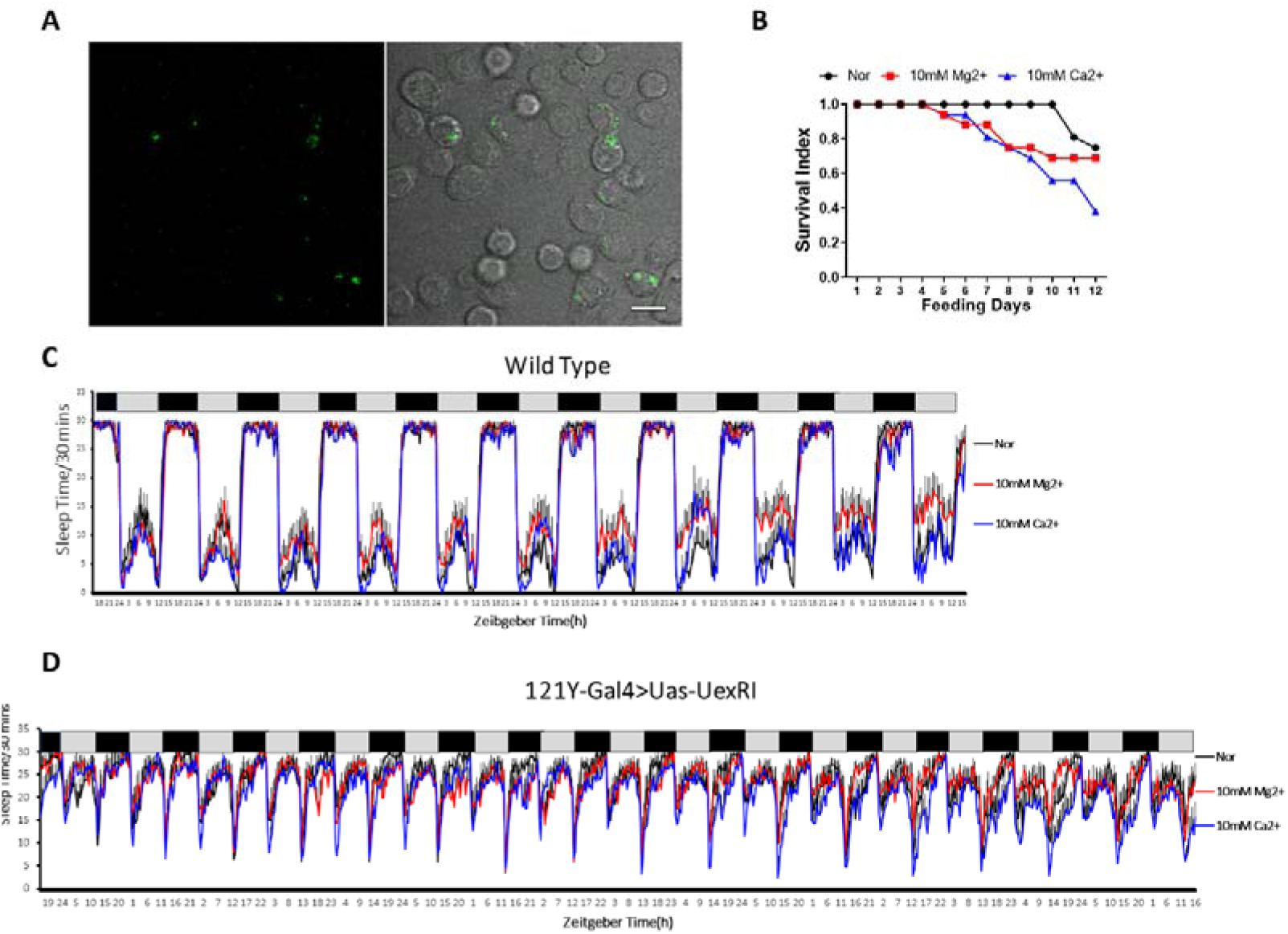
Uex acts to regulate magnesium and Calcium pathway. (A) An illustration of intracellular magnesium green signal. The green magnesium green signal was scattered and aggregated in the cells. Scale bar = 15μm. (B) The mortality rate changed with ion feeding time. Ion feeding increased drosophila mortality, but there was no significant difference between magnesium feeding and calcium feeding. (C) Typical sleep profile of wild typE flies fed with no iron (Black line, n=14), 10mM MgCl_2_(Red line, n=15), and 10mM CaCl_2_(Blue line, n=8). Sleep time was plotted in 30-min bin. The x-coordinate represents zeitgeber time and the y-coordinate represents sleep time every 30 minutes. (D) Typical sleep profile of 121Y-GAL4 > UAS-Uex RNAi flies fed with no iron (Black line, n=14), 10mM MgCl_2_(Red line, n=8), and 10mM CaCl_2_(Blue line, n=14). Sleep time was plotted in 30-min bin. The x-coordinate represents zeitgeber time and the y-coordinate represents sleep time every 30 minutes.

**Figure 6–figure supplement 5.**
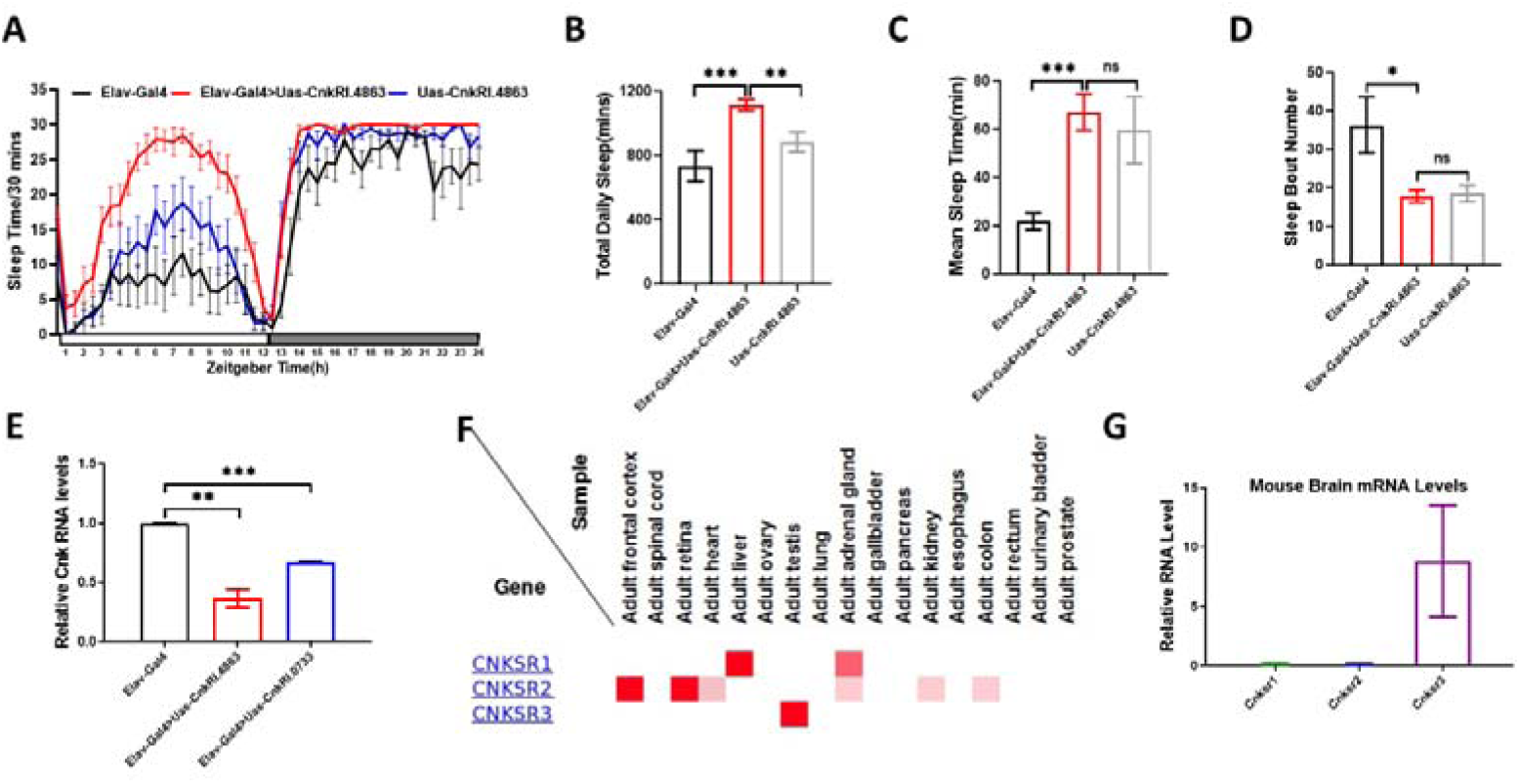
Uex function to regulate CNK/ERK pathway and CNK regulate total daily sleep time. (A)Typical sleep profile of Elav-GAL4 flies (black line, n=16), UAS-Cnk RNAi flies (blue line, n=16) and Elav-GAL4 > UAS-Cnk RNAi 4863 flies (red line, n=16). Sleep time was plotted in 30-min bin. The x-coordinate represents zeitgeber time and the y-coordinate represents sleep time every 30 minutes. (B)The total daily sleep quantification in (A). (C)The mean sleep bout duration quantification in (A). (D)The number of sleep bout quantification in (A). (E) Relative mRNA levels of Cnk in Elav-GAL4 (black column, n=3), Elav-GAL4 > UAS-Cnk RNAi 0733 flies (blue column, n=3) and Elav-GAL4 > UAS-Cnk RNAi 4863 flies (red column, n=3). (F)Tissue expression data of human CNKSRs. The data derived from the Human Protein Atlas database. The deeper the red color, the higher the protein expression. (G)Relative mRNA levels of Cnksr1(black column, n=3), Cnksr2(blue column, n=3), and Cnksr3(red column, n=3) to ribosomal protein L17 in mouse brain.

## Figure legends for Source Data

- Figure 1 Source data1: datasets
- Figure 2 Source data1: datasets
- Figure 3 Source data1: datasets
- Figure 4 Source data1-3: datasets and original gels for Western blot
- Figure 5 Source data1: datasets
- Figure 6 Source data 1-9: datasets, and original gels and blots
- Figure 7 Source data1: datasets
- Figure 1-Supplement 1A-Source data 1-2: datasets and original gels
- Figure 6 - Figure supplement 5 - Source data: datasets

